# Global Nernstian astrocytic depolarization breaks down during local synaptic input

**DOI:** 10.64898/2026.06.23.734112

**Authors:** Ryo J. Nakatani, Erik De Schutter

**Affiliations:** Computational Neuroscience Unit, Okinawa Institute of Science and Technology, Onna-village, Okinawa, Japan

## Abstract

Substantial progress in glial electrophysiology has revealed that astrocytes, which account for half of the cells in the human brain, exhibit membrane potentials that often reflect changes in the extracellular environment. Such responses are mediated by a variety of biochemicals, including potassium and neurotransmitters. Recent advances in voltage imaging have provided new insights into voltage activity in astrocyte peripheries, revealing highly localized depolarization that depends on local presynaptic activity. However, the electrophysiological properties of these isolated peripherals have not been explored due to limitations of spatial and temporal resolution. In this study, we aimed to explore differences in the electrophysiological response between whole-cell stimulation and isolated stimuli at different locations in the cell. Therefore, we constructed an empirical conductance-based NEURON model using a realistic morphology to simultaneously capture both astrocyte processes and soma electrophysiological dynamics. Our results predict a breakdown of the Nernstian behavior of astrocytes when potassium stimuli are localized. Instead, local responses are governed by their conductance ratios. Furthermore, we observe strong capabilities for isolating neurotransmitter responses to specific synaptic inputs, with minimal effect on the astrocyte soma. Our study highlights asymmetrical responses of astrocytic electrophysiology that depend on the spatial scale of stimulation.

**Significance Statement:** Classic astrocyte electrophysiology dictates that the astrocyte membrane potential is strongly coupled to the Nernst potassium reversal potential, which defines the astrocyte’s capacity to buffer potassium and neurotransmitters.

Here, by utilizing a computational conductance-based model, we show that this standard Nernstian relationship breaks down for local synaptic inputs. Our results suggest astrocytes can tune their electrophysiological properties at individual synapses by regulating local conductance ratios. Such capabilities predict that astrocyte function is compartmentalized to the level of individual synapses. These findings provide a framework for designing experiments that directly probe the role of astrocyte buffering at synapses.

## Introduction

Astrocytes, which account for more than half of all cells in the human brain, have generally shown passive responses to classic electrophysiological techniques. However, these non-excitable cells can exhibit dynamic membrane potential responses, often following Nernstian potassium reversal potentials (E_K_). Many studies have explored various conditions that alter astrocyte resting membrane potentials, including changes in extracellular potassium concentrations ([K^+^]*_o_*) or genetic knock-out of specific K^+^channels [1, 2]. These studies reveal that the K^+^ electrode-like response of astrocytes is mediated by various K^+^ channels, which dominantly contribute to astrocyte conductances, with maturation improving astrocyte passivity [3]. These high conductances are supported by the inwardly rectifying potassium 4.1 (Kir 4.1) channels, two-pore domain potassium channels (K2P) and delayed rectifying potassium channels [3–5]. Additionally, many studies in culture have also reported cases in which astrocytes depolarize independently of the Nernst potential. A portion of such cases occurs in culture experiments that observe responses to various neurotransmitters, mediated by both metabotropic and ionotropic receptors. [6–9]. Unfortunately, there are significant discrepancies between culture experiments and *ex vivo* experimental evidence. Many *ex vivo* voltage clamp studies show neurotransmitter currents to be mediated by transporters [10, 11], rather than by ionotropic neurotransmitter receptors [12]. On the other hand, transcriptomic evidence, as well as other molecular reporter models, have suggested functional expression of ionotropic neurotransmitter receptors *ex vivo* and *in vivo* [13–15].

Oftentimes, such contradicting evidence is attributed to differences in expression profiles specific to brain region or astrocyte reactivity [16–18]. However, maybe morphology and local conductances can also explain the observed differences. Electrophysiological techniques are often hampered by space-clamp problems, with morphological attributes significantly contributing to these limitations [19–21]. In cells like astrocytes, with fine branching structures and low input resistance, space-clamp problems are much more prominent. Many technologies have been used to overcome such difficulties, including single-cell dual-patch-clamp, voltage-sensitive dyes, and genetically encoded voltage indicators (GEVIs) [22–24]. For example, single-cell dual-patch-clamping techniques can provide reliable readouts of current-voltage relationships recorded from the soma, but do not necessarily ensure proper voltage-clamping at the periphery. Voltage-sensitive dyes resolve such spatial limitations, but signals are often contaminated with stronger neuronal activity. A recent study using GEVIs specifically resolves this issue by voltage-imaging peripheral signals isolated to astrocyte subcompartments [24]. This study revealed a decoupling between soma and periphery depolarizations, with clear differences in peak depolarization amplitudes detected by GEVIs and somatic patch-clamp recordings.

Therefore, we sought to explore the electrophysiological properties of astrocyte depolarization for peripheral localized microdomains, relative to soma and whole-cell properties. Although many models examining K^+^ dynamics for whole-cell or perisynaptic processes exist [25–28], models that unify the dynamics and interactions between the two are scarce [29–31]. In this study, we constructed an empirical conductance-based NEURON model with a realistic morphology to capture both local and whole-cell astrocyte properties. Using this model, we examined experimental evidence from culture, *ex vivo*, and *in vitro* studies to construct a unifying framework that explains observed classic results and apparent experimental discrepancies. We specifically focused on how the astrocyte response depended on the spatial scales of stimulation and whether this affected depolarization in realistic morphologies.

We found that peripheral astrocytic depolarization triggered by physiological conditions is undetectable somatically, masked by the astrocyte’s high conductance. Our results further suggest two distinct mechanisms of astrocytic depolarization that affect astrocyte membrane potentials in distinct spatial regimes. Our study provides a crucial hypothesis regarding asymmetrical electrophysiological responses to global versus local stimuli and highlights membrane responses that may be specific to synaptic function under physiological conditions.

## Results

### Astrocytes show Nernstian properties for whole-cell potassium manipulations

We constructed an astrocyte potassium-focused whole-cell model to examine the interaction between potassium (K^+^) and the membrane potentials in a realistic morphology. We used a previously reported whole-cell morphology (Fig 1 A; Fig A,B), which contained astrocyte nanoscopic structures recreating characteristics of perisynaptic astrocytic processes (PAPs) measured in electron-microscopy (EM) [29]. The constructed conductance-based model included the major astrocytic K^+^ conductances, such as Kir 4.1, Na/K-ATPase (NKA) and K2P leak channels TREK-1/TWIK-1. These channels were fit to experimental data to recreate empirical electrophysiological properties (Fig S2). The model also incorporated connexin (CX43) conductances at the periphery to recreate gap junctions and astrocyte syncytium properties. Other channels relevant to neurotransmitter mediated astrocytic depolarization, such as glutamate transporters (GLT-1), NMDARs and GABA*_A_*Rs were included in the model for subsequent experiments. The location of PAPs was randomly selected as compartments that were less than 300 nm in length and 500 nm in diameter. The morphological characteristics of the PAP were chosen to match experimental results in Arizono et al. [32]. The PAP morphologies exhibited expected properties such as a higher local input resistance when compared to the soma (Fig A). The model we constructed was specifically designed with a focus on membrane potential and extracellular potassium (K^+^), requiring channel models to be dynamically responsive to changes in the potassium reversal potential (E_K_). As we wanted to examine how channel composition would affect astrocyte [K^+^]*_o_* responsiveness, previously established astrocytic channel models with built-in membrane potential sinks were excluded [33]. The extracellular space (ECS) was implemented to match EM measurements of the extracellular matrix, and calculated K^+^ diffusion based on Fick’s law (detailed derivation in SI text S1).

**Fig 1.**
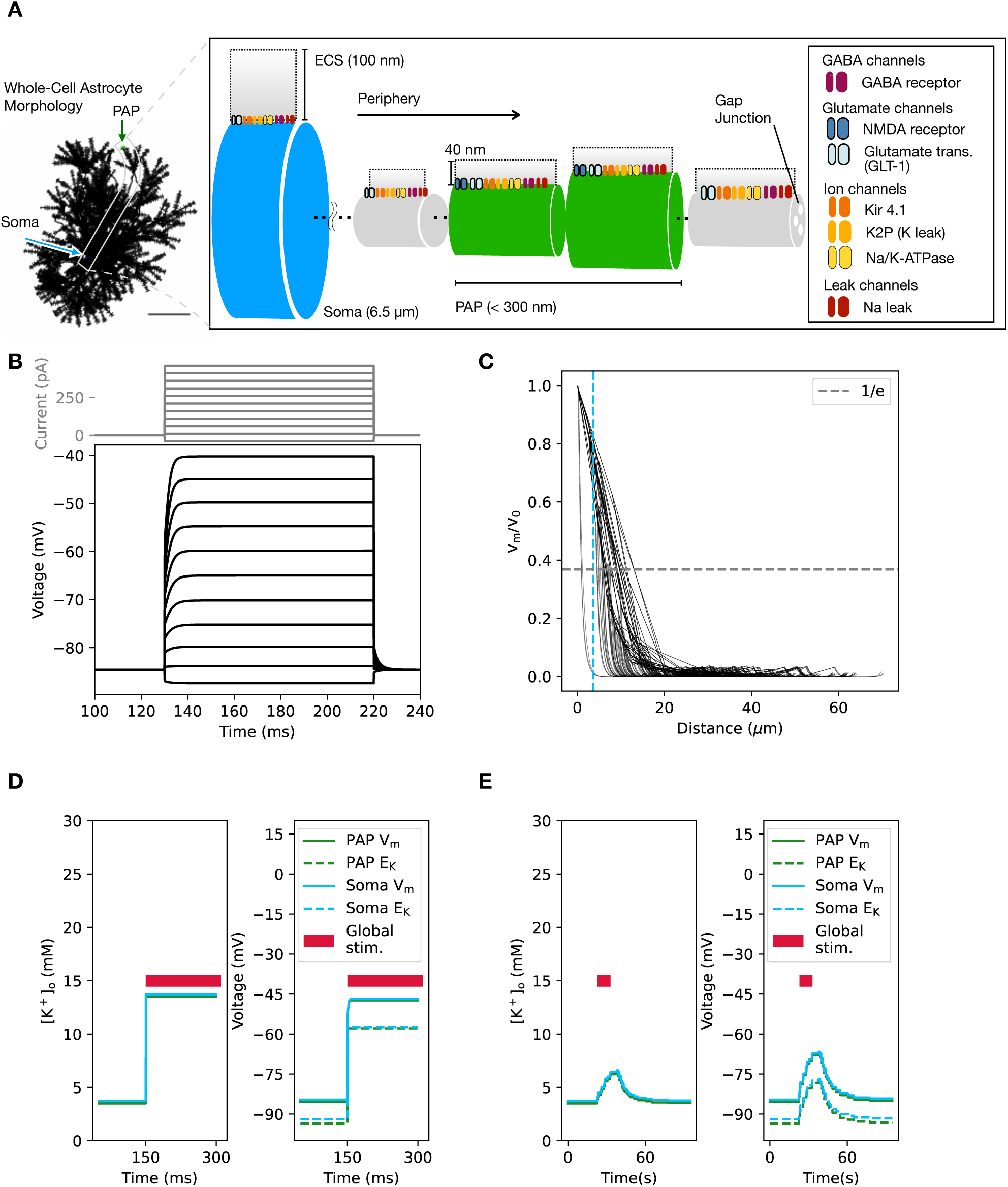
Comparison of the model to whole-cell *in vitro* experiments **(A)** Cartoon depicting our conductance-based model. The astrocyte morphology is that of a previously constructed model [29], with major K^+^ channels Kir 4.1, K2P astrocyte leak channels, Na/K-ATPase and neurotransmitter-related transporters/channels such as GABA*_A_*R, NMDAR, and GLT-1. Equations for Kir 4.1, K2P channels and for GLT-1 were set up to reflect [K^+^]*_o_* changes. Only sections of the model up to 300 nm in length and diameter less than 0.5 µm were considered a PAP candidate. The extracellular compartment was 40 nm in depth to recreate experimental values for PAPs [86 87]. The extracellular compartment for the soma was expanded to 100 nm, assuming the peripherals were more compacted. Extracellular compartments for sections in between were linearly interpolated. As the synaptic receptor distribution was unknown, only PAP sections had synaptic receptors, unless stated otherwise. For all other channels, all distributions were uniform to match densities defined for the PAP. All sections have a sodium leak current to maintain RMP, where current density was calculated analytically (See SI text S9). Each cylinder in the cartoon represents a cable section of the astrocyte model, with blue representing the soma and green representing a PAP. See also Fig S2 and supplementary text S1∽ S2 for justification of the model. **(B)** Soma voltage response to various current injection amplitudes in the model. The current commands are plotted above the figure and range from -40 ∽ 460 pA with 50 pA increments. Model membrane potential responses show typical passive features of astrocytes. **(C)** Somatic voltage attenuation when clamping the astrocyte soma to -60 mV. Space constant for branches ranges from 1∽15 µm, with most voltage attenuating before the periphery. The experimentally measured space constant (*ω* = 3.6 µm) [34] for somatic voltage attenuation is plotted as a vertical blue dotted line for comparison. **(D)** Changes in [K^+^]*_o_* and membrane potential for both soma and PAP during the bath application protocol over time. After bath application of 10 mM [K^+^]*_o_*, the membrane potential changes instantaneously, following the change of the E_K_. **(E)** Changes in [K^+^]*_o_* and membrane potential for both soma and PAP during global application of the *in vivo* protocol. The experimental *in vivo* protocol [2] results in a slow change of [K^+^]*_o_* over a time course of seconds. The membrane potential (peak amplitude -66.6 mV in the soma, -67.3 mV in the PAP) follows changes in the Nernst potential, indicating strong coupling between [K^+^]*_o_* and membrane potential.

Next, to reproduce experimental passive properties in mature protoplasmic astrocytes, we fit the number of Kir 4.1 channels and the passive potassium current density (representing TWIK-1/TREK-1) to experimental results. Our model reproduced the experimentally predicted space constant (*ω*) of 3.6 µm measured by somatic dual patch clamp protocols [34] (Fig S3). The amplitude of Kir 4.1 currents compared to other passive/pump currents was matched to recreate the experimentally predicted 1:1 ratio for potassium currents (I_K_) [22]. With the fitted model, we observed passive responses to current injection at the soma, as reported in classic literature [22] (Fig B). Our results show a generally leaky cell with measured space constants in the range of 1 to 15 µm (Fig C). Previous literature shows that the membrane potential of astrocytes follows that of E_K_ [35]. Bath application of 10 mM [K^+^]*_o_* induced depolarization to -46.87 mV that followed the changes in E_K_ (Fig D). We also observed the effects of gap junctions during these bath application protocols, where inhibition of these channels consistently decreased the K^+^ clearance rate, which matches experimental findings [36] (Fig A,B). To further investigate how experimentally measured *in vivo* global [K^+^]*_o_* changes caused by electrical stimulation could affect depolarization, we next observed the effect of transient K^+^ changes on the membrane potential [2]. Membrane potential strongly reflected [K^+^]*_o_* changes following direct changes to the E_K_, similar to our bath application protocol (Fig E). We also confirm that the general characteristics of astrocyte responses to bath application were not altered by intracellular diffusion (Fig A). Only [K^+^]*_o_* above 10 mM showed stronger coupling to E_K_, which matches the general properties we previously discussed. These results conform with classic astrocyte electrophysiological studies and highlight that astrocyte responses to global stimuli show Nernstian properties.

### Potassium responses of perisynaptic astrocytic processes are governed by local potassium current escape

Next, we sought to identify whether astrocyte membrane potential responses would differ depending on morphology and the spatial confinement of stimuli. We initially examined electrophysiological responses to *in vivo* [K^+^]*_o_* changes solely confined to the PAP or soma. Isolation of the [K^+^]*_o_* changes led to a substantial decrease in depolarization amplitude in comparison to their globally modulated counterparts (Fig A,B vs. Fig E). This was surprising, as the soma in our model contained the highest number of Kir 4.1 channels, suggesting that Kir densities alone did not contribute to depolarization amplitudes (Fig A). Therefore, we next systematically examined membrane potential responses to local [K^+^]*_o_* application by comparing bath application responses with isolated responses at the soma, 4 primary branches, and 25 randomly selected PAPs (Fig C left). All locations showed heterogeneous responses to K^+^, with all exhibiting a loss of Nernstian responses found in the whole-cell regime during bath application (Fig 2 C right). We then confirmed whether this property was altered by adding intracellular diffusion to the model. We observe that the typical [K^+^]*_o_* response of these localized stimuli in the presence of diffusion was indistinguishable for PAPs, primary branches, and soma. This was due to the fast equilibrium of intracellular potassium ([K^+^]*_i_*), as well as the larger volume of intracellular space when compared to the local extracellular compartment (Fig A,B). Therefore, all subsequent simulations disregarded intracellular diffusion to reduce the time required to simulate the model.

**Fig 2.**
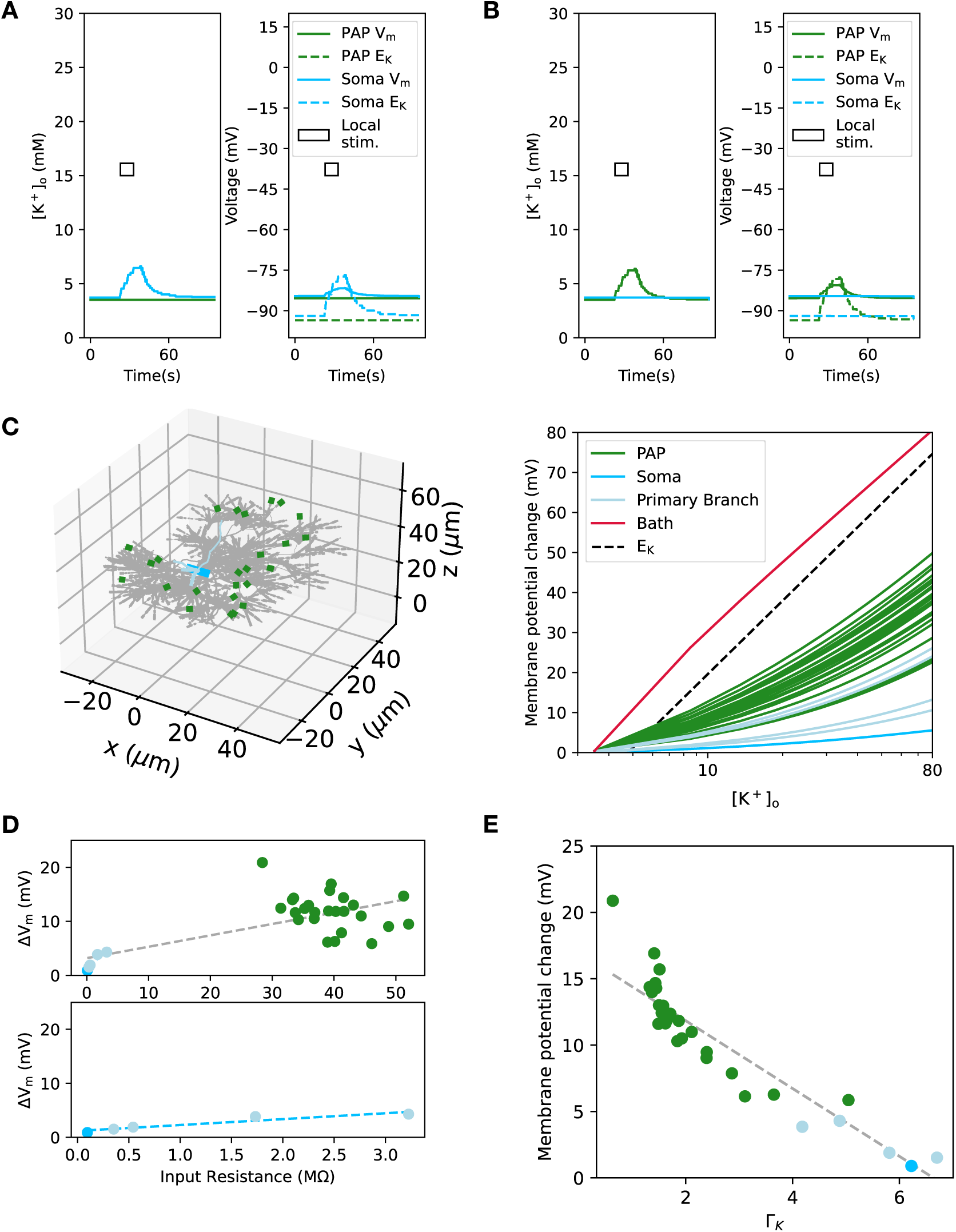
Local stimulation shows non-Nernstian responses **(A,B)** Changes in [K^+^]*_o_* and membrane potential isolated at the A) Soma or B) PAP caused by local application of the *in vivo* protocol. Unlike Fig 1E, the experimental *in vivo* protocol [2] results in a membrane potential that undershoots the Nernst potential for both isolated compartments, with a slightly stronger reduction for somatic response than for PAPs (peak amplitude -81.6 mV in the soma, -80.4 mV in the PAP). **(C)** Left) Location and Right) [K^+^]*_o_*-mediated voltage changes for various local stimuli. Left) Locations for PAPs and primary branches were randomly chosen. The sizes of PAPs are enlarged for visual clarity. Right) All locations show much lower local depolarization responses than in bath conditions that follow the Nernst reversal potential. The [K^+^]*_o_* response is plotted on a log scale for easy comparison between bath conditions and localized conditions. The peak depolarization (ΔV*_m_*) was recorded at the PAP with a single stimulus lasting 0.5 ms. The colors indicate the category for location (Soma, Primary branch, PAP or Bath), corresponding to the legend within the figure. **(D)** Comparison between peak membrane potential changes (ΔV*_m_*) and input resistance. Peak potentials were measured as the response to a local [K^+^]*_o_* change of 10 mM. All data points had a weak correlation (Pearson’s correlation p-value: 4.7 × 10*^→^*^7^ *R*^2^ = 0.54). Data points for soma and primary branch had a stronger correlation (bottom plot; Pearson’s correlation p-value: 1.5 × 10*^→^*^2^ *R*^2^ = 0.88), which was lost in the PAP-specific data. **(E)** Comparison between peak membrane potential changes and local current escape, expressed as Γ*_K_*. Γ*_K_* represents conductance ratios between axial conductance per cross-sectional area (*g_a_*) and specific K^+^ membrane conductance (*g_k_*). 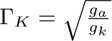 was obtained from a systematic search for an intuitive parameter (details highlighted in methods). All data points had a strong correlation (Pearson’s correlation p-value: 3.0 × 10*^→^*^12^ *R*^2^ = 0.83).

To better understand the local responses, we initially investigated if they were ohmic-dominant. Therefore, we compared the depolarization amplitudes observed with a 10 mM change in [K^+^]*_o_* to values of measured local input resistance. Interestingly, although the soma and primary branches were correlated with input resistance, there was only a weak correlation between all responses and input resistance, which disappeared upon examining correlations between PAPs and input resistance (Fig 2 D; total *R*^2^ = 0.54, soma and primary branch only *R*^2^ = 0.88). This suggests that the local [K^+^]*_o_* mediated depolarizations were not ohmic-dominated and that there was stronger heterogeneity at the PAPs. Given that input resistance alone could not explain [K^+^]*_o_*mediated responses, we next focused on electrophysiological parameters that were intuitive and, ideally, experimentally measurable. After various comparisons of electrophysiological parameters (data not shown), we found that the ratio of axial conductance per cross-sectional area (*g_a_*) to specific potassium membrane conductance (*g_k_*) strongly explained variations in [K^+^]*_o_* responses (details for derivation in methods). Briefly, this parameter can be obtained using two experiments: first, measuring the local input resistance; and second, measuring the [K^+^]*_o_* change required to generate a current of arbitrary amplitude. The results from these protocols were then used to obtain the ratio of effective axial conductance to effective potassium conductance. We further found that the square root of this ratio was the best-fitting (Fig E; *R*^2^ = 0.83). We defined this parameter (Г*_K_*) as:

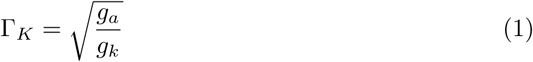

Intuitively, this parameter represents the local capability to regulate the escape of I_K_ at that location, with large Г*_K_* resulting in greater escape to neighboring compartments. The escaping current mostly disappears by the neighboring membrane conductances, as expected from the small space constants of the cell. We observed that typical PAPs with minimal surface area exhibited large variations in neighboring structures, resulting in substantial variations of Г*_K_*. With constricting neighbors, we saw lower values of axial conductance (e.g. Г*_K_*) and higher [K^+^]*_o_* responses. Furthermore, the PAP with the highest membrane response, nearing the Nernst potential, had the lowest Г*_K_*. On the other hand, primary branches and the soma tended to have large surface areas and multiple neighbors due to branching points. This explains the high value of axial conductance (e.g. Г*_K_*) and lower [K^+^]*_o_* responses. Although the parameter seems similar to the space constant in passive cables, it differs critically, as it is only a unitless ratio of conductances. The strong dependence on Г*_K_* highlights the importance of axial conductances for local depolarizations, blocking the Nernstian properties observed in bath application.

### Presynaptically driven astrocyte depolarization requires excessive localized potassium concentrations

So far, our model’s results suggest that astrocytes exhibit distinct electrophysiological dynamics that depend on stimulus localization. We therefore next compared our results to the experimental literature in physiological contexts. Results reflecting the electrophysiological properties of isolated stimuli are limited, except for those using genetically encoded voltage indicators to assess neuronal axon-driven responses of astrocytes in somatosensory cortex brain slices [24]. In this experiment, astrocytes displayed isolated 20 mV depolarizations in response to stimulating ascending axons under dl-2-amino-5-phosphonopentanoic acid (APV) and 6,7-dinitroquinoxaline-2,3-dione (DNQX). Such results would suggest large depolarizations confined to PAPs, independent of postsynaptic components. Typically, [K^+^]*_o_* contributions to the synaptic cleft are from postsynaptic neuronal *ε*-amino-3-hydroxy-5-methyl-4-isoxazolepropionic acid receptor (AMPAR) and NMDAR activity [37]. However, the depolarization amplitudes in [24] were observed in the presence of neuronal synaptic receptor blockers, indicating mechanisms of depolarization dependent on presynaptic effects. The simulation protocols had a focus on the [K^+^]*_o_* change induced by a single action potential (AP) during neuronal repolarization, typically lasting *→* 0.5 ms. The [K^+^]*_o_* amplitude was estimated using values from previous computational models of [K^+^]*_o_* in the synaptic cleft [33, 38]. Therefore, the simulation protocol used simplified action potentials (sAP; 0.5 mM increase of [K^+^]*_o_* for 0.5 ms) for 10 pulses at 100 Hz, matching the experimental protocol. This sAP stimulation resulted in minimal depolarizations that did not match experimental values, as there was no [K^+^]*_o_* build-up between APs within the ECS of the PAP (representative trace Fig A). The lack of accumulation of [K^+^]*_o_* in our model was due to the quick removal of K^+^, mainly by diffusional leak of K^+^ from the small ECS (Fig S6 A). The difficulty in accumulation was further verified by an analytical model of Kir 4.1-mediated K^+^ uptake with no loss by diffusion, suggesting a half-life of less than 1 ms for uptake of 20 mM K^+^ in realistic ECS sizes (Fig B,C). These results suggest that local diffusion and Kir 4.1 uptake both strongly inhibit K^+^ accumulation due to local [K^+^]*_o_*elevations within a realistic ECS.

To verify that this effect is robust regardless of channel composition, we next altered channel densities for Na/K-ATPase and glutamate transporters (GLT-1), to examine how K^+^ uptake and depolarization by these channels would differ. Interestingly, altering NKA densities and GLT-1 had minimal effect, even with the application of glutamate in the ECS (Fig A,B). These results suggest that K^+^ dynamics are mainly driven by Kir 4.1, and other channel composition contributes less to the membrane potential than K^+^ amplitude. Such results match experimental findings, where 50 ∽ 80 % of I_K_ were derived from Kir 4.1 channels [22, 39].

Given our previous results highlighting the difficulty of depolarizing astrocytes with local physiological stimuli, we next set out to explore various AP scenarios that could effectively depolarize astrocytes with isolated synaptic input. First, we considered alteration in the amplitude of the 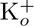 stimuli. To reach experimentally observed values, large amplitudes of 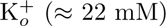 were necessary for considerable depolarization (representative trace Fig 3 B). The K^+^ requirement of 22 mM for depolarization amplitudes of 20 mV was the median for all observed PAPs, which had a range of 17.5 ∽ 62.7 mM (Fig 3 C, Fig S8).

**Fig 3.**
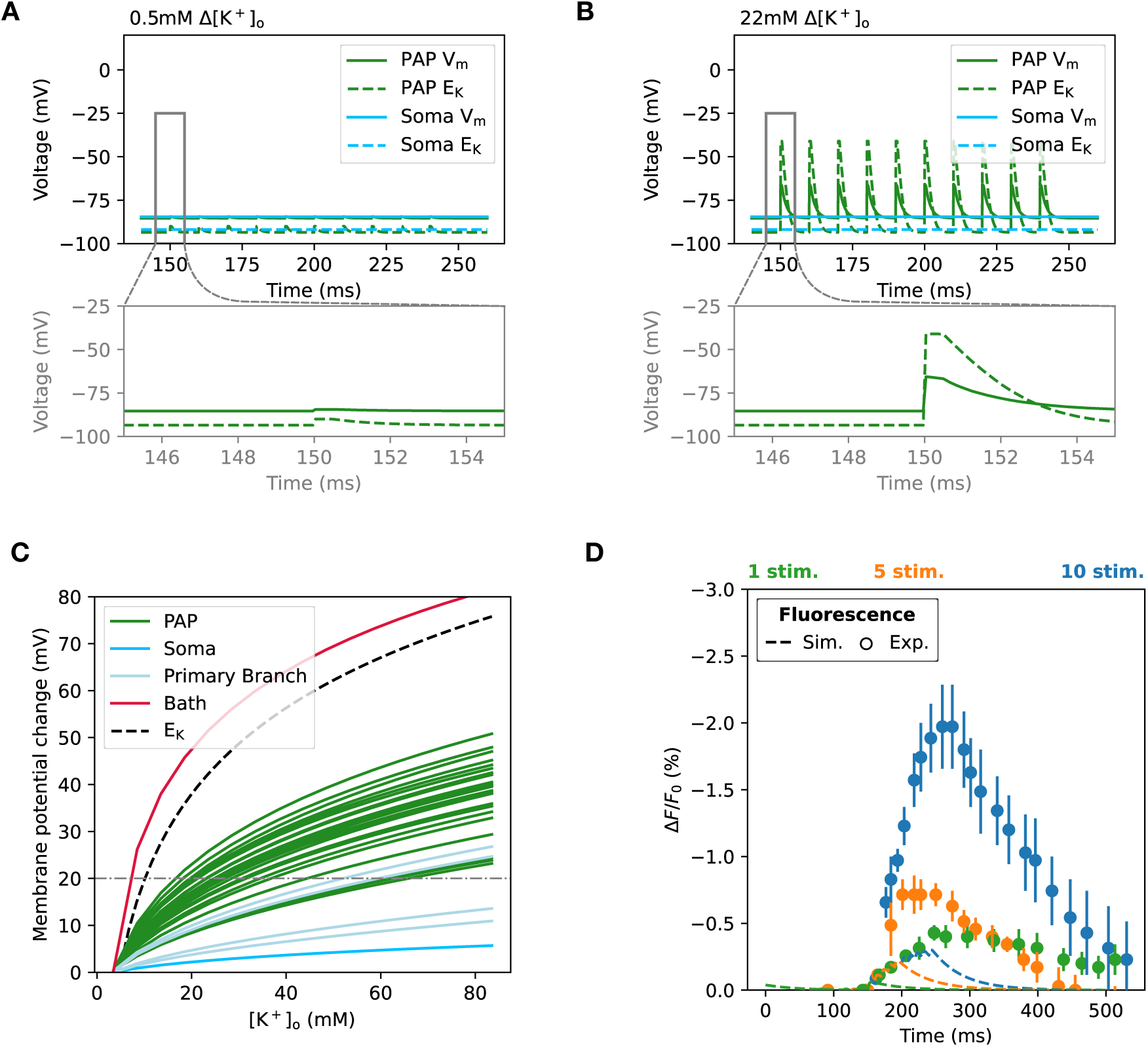
Presynaptically released potassium-driven depolarization shows diminished depolarization **(A,B)** Depolarization of model by A) 0.5 mM or B) 22 mM [K^+^]*_o_* changes using the amplitude modified sAP protocol applied to the PAP. The sAP had a [K^+^]*_o_* stimulus with a duration of 0.5 ms. Top) Membrane potential changes over time in the PAP and soma induced by the modified sAP. Bottom) An inset showing changes in membrane potential for a single stimulus. Simulations were done with 2630 Kir 4.1 channels, which was the default value fit to match experimental results [22 34]. **(C)** [K^+^]*_o_* responses for all isolated locations replotted to showcase [K^+^]*_o_* needed for a 20 mV response. The plot contains the same data as Fig 2 C, but the x-axis is no longer on a log scale. **(D)** Model fitting to experimental results observing depolarization [24] mediated by all K^+^ channels, and GLT-1 in our astrocyte model. The y-axis shows changes in the fluorescent voltage indicator (ArcLight; Г*F/F*_0_) for both simulation and experiment. We simulate fluorescent voltage indicator dynamics by matching time delay dynamics observed by Milosevic et al. [40]. The results are poor in comparison to experimental values, when stimuli were well isolated at the synapse (SSE: 32.1). Note the small difference between simulated peak amplitudes for 5 or 10 stimuli compared to the large difference in the experimental data. Fitting was achieved by using Δ[K^+^]*_o_*=22 mM, 2630 Kir channels in PAP.

We next compared our model with previously conducted ArcLight experiments [24], which have a measured calibration curve converting fluorescent changes to membrane potential changes. By implementing ordinary differential equations that match the time dynamics of ArcLight within our simulations, we acquired pseudo-fluorescence changes reflecting our model’s membrane potential responses [40] (Fig A). In this comparison we utilized the previous 22 mM [K^+^]*_o_* stimuli that exhibited 20 mV depolarization. As a result, we show that with this stimulus experimental traces could not be matched (Fig 3 D, D, Fig S10; Single synapse model SSE: 32.1). As mentioned before, these depolarizations were much smaller than expected Nernstian values. These predicted depolarizations were consistently diminished compared to the actual depolarization amplitudes, further implying the limited capabilities of fluorescent indicators for detecting fast transient responses (Fig S9 B). These results indicate the difficulty of achieving depolarizations with presynaptically increases of [K^+^]*_o_*, as responses are diminished by local current escape and rapid diffusion.

### Responses of perisynaptic astrocyte processes are more affected by spillover than by channel composition

Our previous results highlight the difficulty of local [K^+^]*_o_*-mediated depolarization compared with conventional global application. Based on this observation, we next asked how depolarization dynamics would be altered by clustered synaptic inputs rather than single presynaptic stimuli. Therefore, we examined how increasing the ‘affected PAP lengths’ (i.e. the spatial range of perisynaptic stimuli) can affect astrocyte membrane potentials. We suspected that such a response would lie between the global Nernstian and the local properties we previously observed. As a caveat of our modeling, we did not consider 3-dimensional aspects of extracellular diffusion. To systematically observe how changes in localized activation would alter depolarization dynamics, we increased these lengths from typical PAP lengths of 0.3 µm to 10 µm, which are lengths comparable to the soma diameter. By spatially extending the sAP stimuli of both K^+^ and glutamate, we find that lengths above 2 µm have depolarizations of over 2 mV (Fig 4 A). Our model compares well with experimentally estimated values of glutamate transporter-mediated depolarization of 3-4 mV in PAPs [24]. These results suggest that depolarization of astrocytes via GLT-1 can be triggered only by strong, repeated stimuli that produce largely synchronized K^+^/glutamate release and spillover.

**Fig 4.**
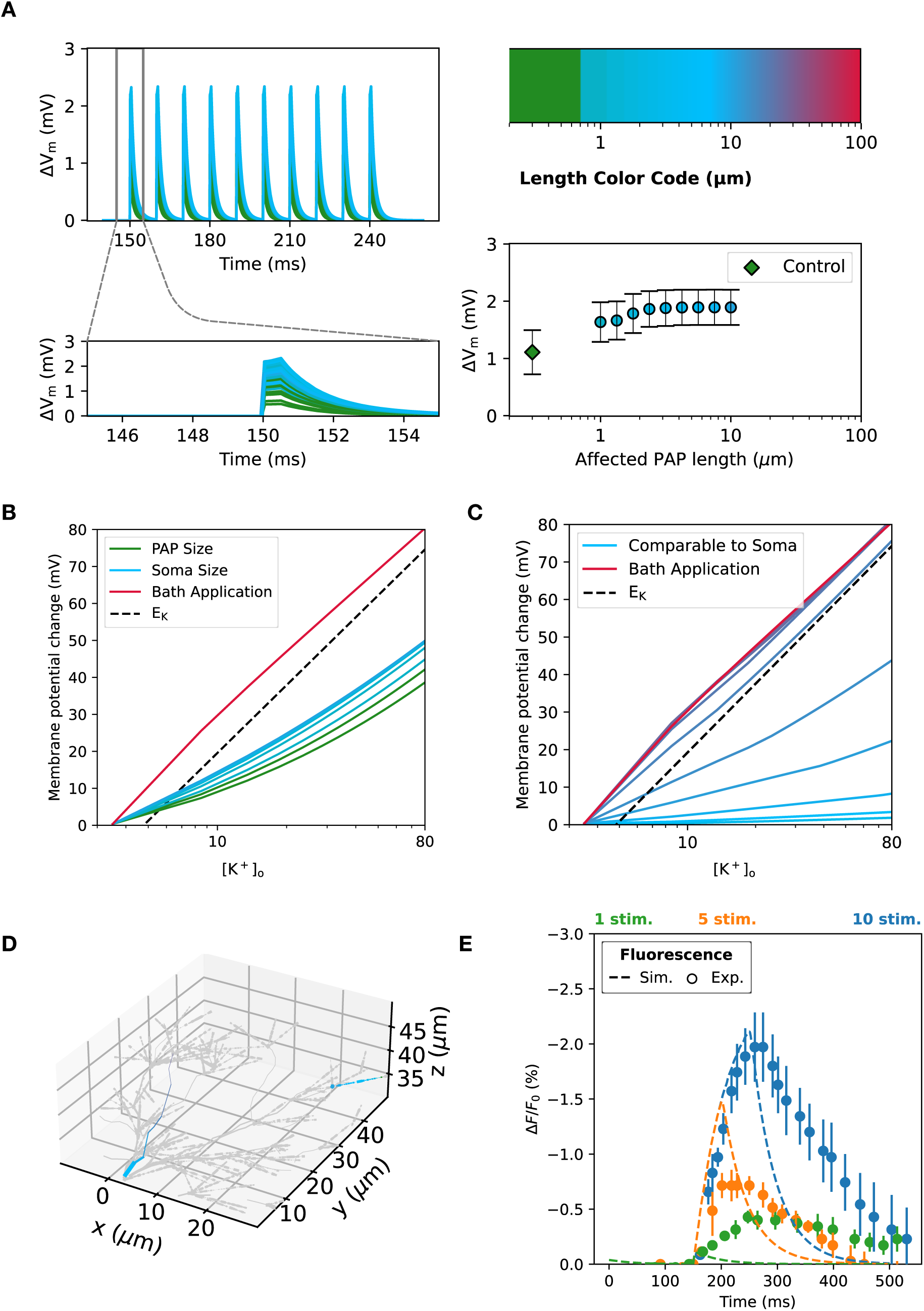
Local spillover enhances depolarization amplitudes **(A)** Left) Currents of each individual altered PAP length are indicated by color in the legend in the top-right (*n* = 10 for each length). Insets on the bottom show a single response to [K^+^]*_o_* (0.5 mM) and glutamate change (1 mM). Saturation of depolarization amplitudes can be observed within the observed ranges. Right bottom) Plots of peak membrane potential changes against PAP lengths with marker colors matching the legend in the top-right. The diamond shown in the plot and within-plot legend indicates the control condition (0.3 µm) of the affected PAP length. Longer lengths than the physiological condition are all spillover conditions, marked with a circle. **(B,C,D)** B,C) Response to [K^+^]*_o_* for spatially extended stimulus conditions (semi-log scale), for locations indicated in D. The color code reflects the spatially extended length marked in A. B) [K^+^]*_o_* response for extended affected PAP lengths, reflecting clustered synaptic input. Changes in PAP lengths are shown in the right branch of D, matching the color code of A. Red is the response for bath application. C) [K^+^]*_o_* response for extended affected PAP length, reflecting a larger spatially extended stimulus. These stimuli are analogous to effective ranges of puff applications on the primary branch (20 ∽40 µm). Changes in PAP sizes are shown in the left branch of D, matching the color code of A. D) The up close morphology of selected locations for B and C. **(E)** Model fitting to experimental results observing depolarization [24] mediated by all K^+^ channels, and GLT-1 in our astrocyte model. The y-axis shows changes in the fluorescence voltage indicator (ArcLight; Г*F/F*_0_) for both simulation and experiment. Slowing down [K^+^]*_o_* dynamics in the synaptic cleft, recreating situations that result in global [K^+^]*_o_* accumulation, decreases SSE (SSE: 11.2). The simulated peak amplitudes now match the experimental data for the response to 10 stimuli, but the fit for 5 stimuli overshoots it. Fitting was achieved by using the same [K^+^]*_o_* input as Fig 3E but altering K^+^ and glutamate decay dynamics in the synaptic cleft (Г[K^+^]*_o_*=22 mM, 2630 Kir channels in PAP, slowing of K^+^ dynamics by 603-fold, glutamate dynamics by 1.08; glutamate time-dynamics had minimal contribution).

We next observed how manipulations to [K^+^]*_o_* in the sAP protocol would affect peak depolarization. The results show a gradual increase in the peak depolarization response, which slightly approached Nernstian values (Fig 4 B, D right section). Since the previous modifications only observed clustered synaptic inputs at peripheral branches, we next examined how larger stimuli near the soma, such as localized puff applications, would alter the depolarization response. By focusing on confocally visible primary branches, we observed that localized [K^+^]*_o_* responses on these branches were significantly increased, with full branch activation triggering Nernstian responses in the astrocyte (Fig 4 C, D left section). These results highlight the critical differences of astrocytic responses between isolated and spillover conditions, with increased lengths of activation contributing to astrocyte responses that approach Nernstian properties.

Using this aspect of increased activation of the model, we fit our model to the ArcLight experiments [24]. Considering that these experimentally observed depolarizations would also induce glutamate release, we also activated GLT-1. Altering affected PAP lengths and slowing synaptic cleft dynamics to represent spillover dynamics yielded better-fitting results than our previous model (Fig 4 E, Fig S10; Spillover model SSE: 11.2). Furthermore, some of the fluorescent traces for the spillover model matched only the initial amplitude, not the decay, of the experimental data. The model peak amplitudes also increased sublinearly with number of stimuli, while the relation for experimental results was supralinear. Additionally, our model shows that the predicted voltages from the fluorescent indicator were strongly decoupled from actual membrane potential peaks, highlighting the limitations of measuring voltages by ArcLight during complex astrocyte responses (Fig S9 C). These results suggest that, with K^+^ mechanisms and GLT-1, depolarization can be achieved only under high-frequency stimulation in near-pathological extracellular conditions and may still require other slow components to match the experimentally observed decay.

### Localization of neurotransmitter channels strongly affects observable somatic voltage-clamp currents

In many pathological cases where neurons are hyperactive, astrocytes have been observed to depolarize along with brain-wide increases in extracellular glutamate and [K^+^]*_o_* [41]. Our previous results in fitting local astrocytic depolarization (Fig 3 D, Fig 4 E) highlighted the impact of [K^+^]*_o_* accumulation over extended distances in the ECS. At the same time, it also reveals the lack of a slow-decay component in astrocytic depolarization driven by Kir 4.1, GLT-1, and NKA during isolated depolarization events at the PAP. Therefore, we questioned whether these channels and [K^+^]*_o_* are the only causes of astrocytic depolarization and explored the effects of additional mechanisms known to depolarize astrocytes. Previous experimental results show that sub-compartments of astrocyte activity are coupled to neuronal activity, suggesting that astrocytes can respond to neurotransmitters [15, 42, 43]. Other classic electrophysiological studies of astrocyte neurotransmitter responses in cultures have also suggested neurotransmitter-mediated depolarization [6–9]. However, there is also a strong lack of evidence in *ex vivo* studies, with many cases showing minimal to no neurotransmitter-mediated currents. Often these differences between *in vitro* and *ex vivo* are attributed to astrocyte reactivity induced in cultures, which would strongly alter protein expression in astrocytes [44, 45]. We investigated whether this is the only possible explanation for the discrepancy between experimental observations.

One key difference between *in vitro* and *in vivo* is the maturation of astrocyte morphology. Therefore, we sought to examine the relationship between synaptic transmitter localization and morphology and somatically observable currents. Within our experiment, we placed neurotransmitter-related channels, such as NMDARs (N2C), and GABA*_A_*Rs, along with our previous GLT-1 channels [10, 46–48]. These channels were chosen as they have largely independent electrophysiological properties, and have strong evidence of functional expression in astrocytes of the somatosensory cortex in physiological conditions [14, 15]. Moreover, we focused on ionotropic receptors, as they contribute on a much faster timescale than metabotropic receptors, which is necessary for the rapid accumulation of depolarization during short pulse trains observed in experiments [24].

Evidence suggests that the majority of NMDARs are expressed in the periphery [13, 14, 49], while GLT-1 expression is throughout the whole cell [46, 48]. On the contrary, GABA*_A_*R expression levels depend on brain region, with cortical astrocytes thought to have enriched expression at fine processes, but lacking definitive evidence [50]. Therefore, we localized the channels to their respective domains within our model by excluding synaptic receptors from the primary branch and soma. The models for GABA*_A_*R and NMDARs reflected astrocyte physiology, such as higher reversal potential for GABA*_A_*Rs in astrocytes, and weaker susceptibility to Mg^2+^ in NMDARs. We then split the whole cell into 10 different shells based on distance from the soma (Fig 5 A), and randomly placed 40 synapses in each shell, which was necessary to fit experimentally measured GLT-1 amplitudes observed in the somatosensory cortex [51]. By examining the somatic voltage-clamp electrode current response to activation of synapses in each shell, we saw a reduction in the ratio between somatically observed electrode current and the sum of actual GLT-1 induced currents with distance from the soma (Fig 5 B). Such a phenomenon is attributed to space-clamp problems [19–21], expected in cells with complex morphology and high membrane conductance [22]. Although channels, such as GLT-1, with high expression throughout the cell can be detected in these situations, their current amplitudes were strongly underestimated (Fig 5 B). We then compared the somatically observable voltage-clamp current of each neurotransmitter channel, upon synaptic activation of all synapses in all shells (Fig 5 C). A comparison of synaptically induced current to somatic measurement for each shell reveals that for NMDAR and GABA*_A_*R placed on peripheral branches, there are no observable somatic currents regardless of strong synaptic input (Fig 5 C). We additionally examined effects of sparse global placement of GABA*_A_*Rs and global puff application of GABA in our model, similar to observed experimental protocols in hippocampal astrocytes, and saw our results were similar to experimentally observed somatic changes in these *ex vivo* experiments [52] (Fig S11). This supports the concept that most astrocyte currents in the periphery will be masked during somatic voltage clamp by the K^+^ conductance dominating the cell, excluding transporter channels such as GLT-1, which have abundant expression throughout the cell [53]. These results suggest that the lack of experimentally observable synaptic receptor currents *ex vivo* may be due to the localization of these receptors in synaptically dense areas, often far from the soma.

**Fig 5.**
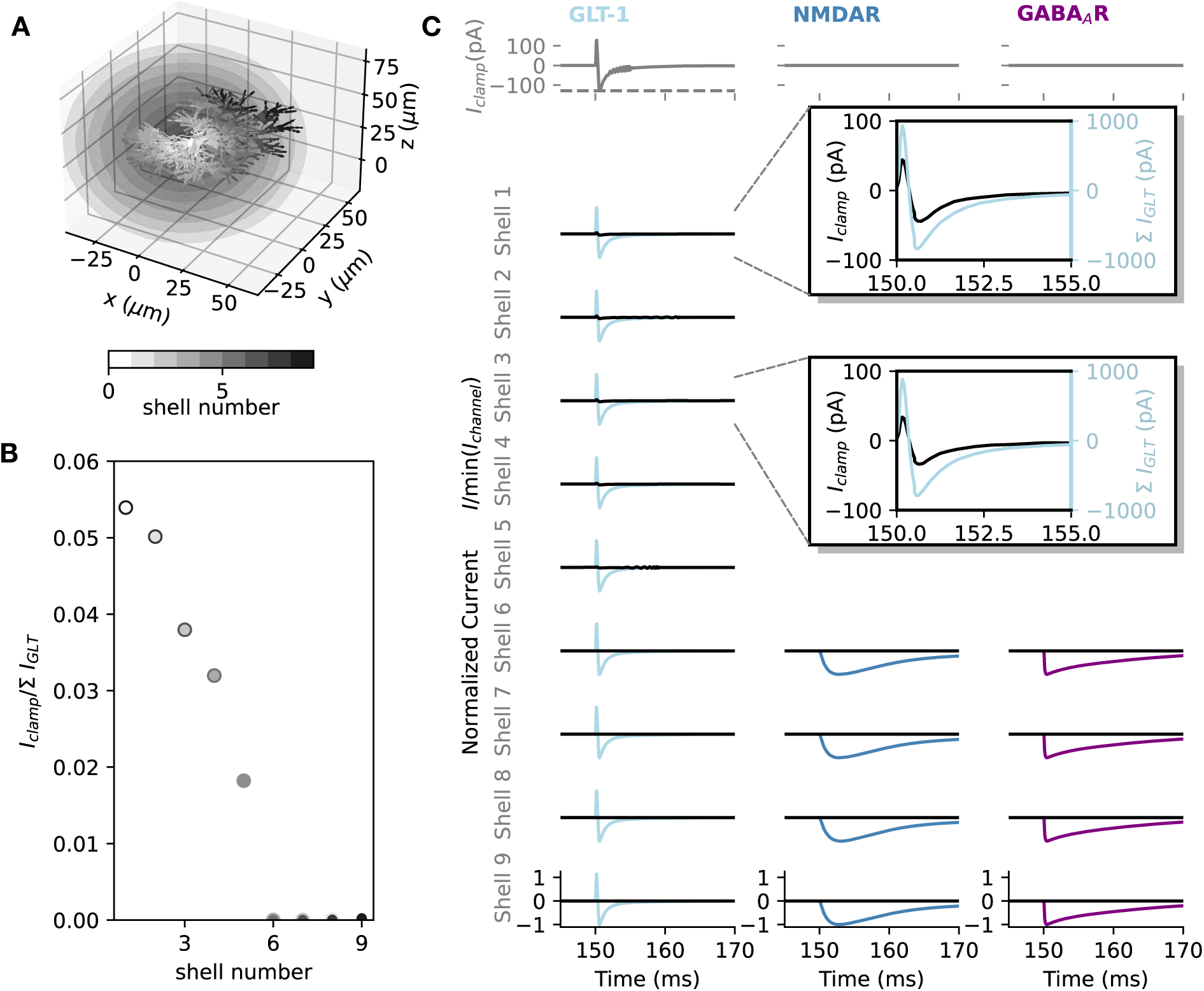
Effects of distance on somatically observable voltage-clamp currents **(A)** Morphology of the astrocyte used in the model. The morphology was split into 10 shells based on the distance from the soma and color-coded as shown in the scale-bar. Shell 1 ∽ 5 contains the soma to the primary branches of astrocyte morphology, and the following shells are mainly composed of the branchlets and PAPs of the astrocyte model. **(B)** Ratio between total GLT-1 currents (# *I*_GLT_) per shell and observable voltage-clamp current (*I*_clamp_) at the soma. The ratio of observable current and GLT-1 current decays as shells reach more peripheral sections. Almost no current is observable after shell 5, which is when all primary branches have been excluded. **(C)** Relationship of measured voltage-clamp currents and synaptic currents plotted over time. The top row indicates the somatic voltage clamp current recorded upon stimulation of all available synapses in all shells, showing a complete lack of observable GABA*_A_*R or NMDAR induced currents. Experimentally observed amplitude of GLT-1 induced current [51] is plotted as a dotted horizontal line in the top-left trace. Subsequent rows show the breakdown of normalized total synaptic current (*I/* min(*I_channel_*)) and corresponding somatic voltage clamp current per shell for each synaptic channel type. The somatic current becomes smaller as shells reach peripheral compartments. Synaptic receptors were placed only in compartments that excluded the primary branch and soma (shells 6 to 9). Shell 0 was excluded because it contains only the soma and is assumed to have no direct synaptic contact. Two insets show the actual current time courses and values, where the black curve and left y-axis correspond to the observed somatic current (*I*_clamp_). The light blue curve and right y-axis correspond to the total GLT-1 current (# *I*_GLT_). The light blue axis is scaled by 10 times for better visual comparison to the black axis.

### Neurotransmitters can mediate focal depolarization independent of potassium

Given our previous results suggesting that neurotransmitter receptors may be present at electrophysiologically undetectable locations, we next sought to understand how they might impact PAP depolarization. We therefore examined how GABA*_A_*Rs could hypothetically contribute to local astrocyte depolarization and questioned what depolarization amplitudes could be observed in astrocyte sub-compartments parallel to synapses. We examined the peak depolarization in the PAP by adding 1.5 mM of GABA after the [K^+^]*_o_* stimulus to recreate neurotransmitter release [54]. Using the same 10 pulse 100 Hz stimulus of sAPs, we saw it was possible to obtain depolarizations observed in experiments. Alterations of GABA*_A_*R densities strongly affected depolarization amplitudes (Fig 6 A). Although the depolarization was sustained for longer than that of Kir 4.1 only, voltage attenuation maintained the isolation of PAP voltage dynamics (Fig 6 B). GABA*_A_*Rs also produced depolarization that did not require large [K^+^]*_o_* or longer PAP lengths, which were key factors for astrocyte depolarization in our previous results. The contradiction to typical neuronal GABA*_A_*R contributions is due to the different intracellular chloride concentration of astrocytes, leading to depolarizing GABA*_A_*R currents [47, 55]. On the other hand, during the depolarization of the PAP by GABA, the membrane potential rose higher than E_K_, resulting in an efflux of K^+^ from the cell (Fig 6 C). To further analyze how the GABA*_A_*R channel contributes to depolarization when compared to Kir 4.1, we examined the phase plane of [K^+^]*_o_* against membrane potential. We revealed that more activated GABA*_A_*Rs induce larger depolarization amplitudes, while larger Kir 4.1 densities create stronger sinks to E_K_ (Fig 6 D). We next fit our model again to experiments [24], which yielded better results than the Kir 4.1/GLT-1 model without spillover, although the 10 stimuli response largely undershoots the expected experimental peak (Fig 6 E, Fig S10; SSE:17.5).

**Fig 6.**
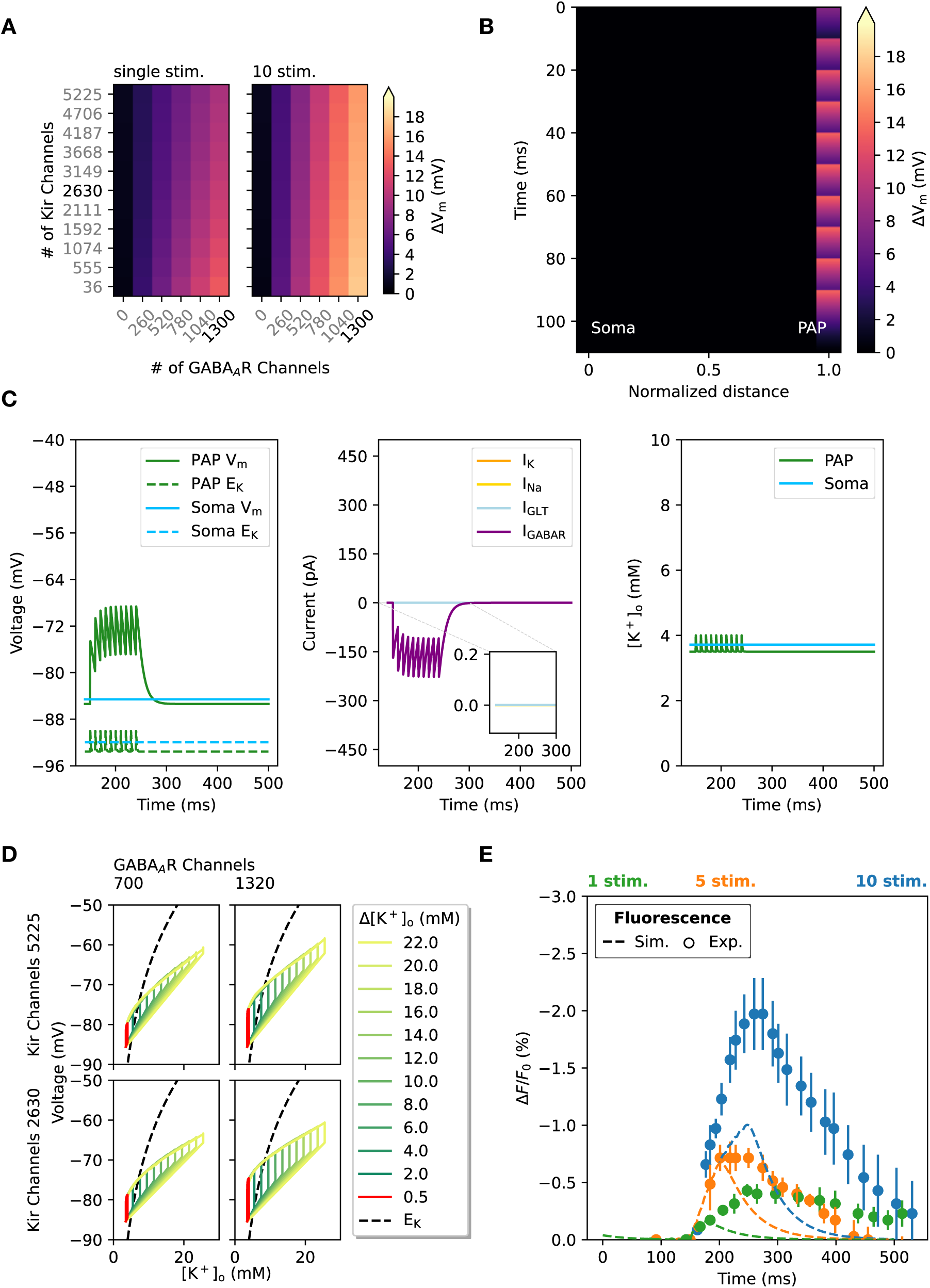
GABA*_A_*R driven depolarization **(A)** Heatmaps showing typical depolarization response of PAPs with various counts of Kir 4.1 channels and GABA*_A_*R. The peak depolarization recorded at the PAP was compared for both Left) single and Right) 10 stimuli of 0.5 ms. All stimuli were followed by a 1.5 mM transient increase of GABA. The colors indicate the peak membrane potential change (ΔV*_m_*) for each condition, corresponding to the color scale within the figure. The black text labels indicate control conditions, while all gray values are models with modified channel numbers. **(B)** The voltage attenuation along the branch from soma to PAP over time, showing a complete lack of depolarization away from the PAP. Distance was normalized to the distance between the PAP far end and the soma. There are 1320 GABA*_A_*R channels and 2630 Kir 4.1 channels to match control conditions used in panel A, C and E. **(C)** Left) Voltage (V*_m_*) along with reversal potential (*E*_K_) for PAP and soma, Middle) currents in the PAP (*I*_K_*, I*_Na_*, I*_GABAR_*, I*_GLT_), Right) [K^+^]*_o_* within the PAP section and soma. Middle plot inset shows all currents other than *I*_GABAR_ to show minimal contribution by these currents, as GLT-1 is not activated by GABA. **(D)** Phase planes of [K^+^]*_o_* vs. voltage for respective combination Kir 4.1 and GABA*_A_*R counts. Traces for phases with different amplitudes of [K^+^]*_o_* stimuli are color-coded and labeled in the legend. Each panel contains four figures with the corresponding conditions labeled for each column and row. All phase planes show the model responses to a single stimulus. The control condition for the relative [K^+^]*_o_* increase we have used for typical sAPs is shown in red. The reversal potential calculated from voltage and [K^+^]*_o_* is plotted as a broken line in black. **(E)** Model fitting to experimental results observing peripheral depolarization in astrocytes [24]. The y-axis shows changes in the fluorescent voltage indicator (ArcLight;Г*F/F*_0_) for both simulation and experiment. Fitting was achieved by using channel counts as a free parameter (Fitted result: 1320 GABA*_A_*R). The fitting results are better in comparison with synaptically confined [K^+^]*_o_* inputs (SSE:17.5), despite a strong undershoot of the 10 stimuli simulated response.

Although GABA-mediated depolarization suggests astrocyte membrane responses to inhibitory neurons, it was logical to ask whether this response was selective to inhibitory synapses. Recent studies have shown behavioral changes induced by knockdown of cortical somatosensory astrocyte NMDARs, suggesting that these channels are functionally expressed [15], with direct implications for astrocyte calcium and function. We examined the peak depolarization in the PAP by using the same stimulation protocol as GABA*_A_*R depolarization, replacing GABA with glutamate release of 1 mM. As a result, we found that NMDAR-mediated depolarization replicates our GABA*_A_*R results, with a strong dependence on NMDAR channel numbers, and voltage isolation from the soma (Fig 7 A-D). However, it is important to note a few key differences when comparing to GABA*_A_*R-mediated depolarization (Fig 7 A-D). Firstly, NMDAR depolarization seemed to have larger accumulatory effects, which are similar to experimental results [24], as NMDARs have longer channel open times than GABA*_A_*Rs (Fig 7 C). Interestingly, single channel conductance was only slightly different between NMDAR (33 pS) and GABA*_A_*R (28 pS) [56, 57], suggesting the open channel times are more critical in contributing to PAP depolarization. Secondly, the change in channel open times also affected the decay time-course of depolarization and [K^+^]*_o_* (Fig 7 C). Lastly, our model shows that depolarization via NMDAR can be done with a smaller number of channels (344 channels) compared to that of GABA*_A_*R (1320 channels) (Fig 7 C vs. Fig 6 C We therefore suggest that NMDAR channels that are expressed at a much lower density than Kir 4.1 channels, GLT-1, and GABA*_A_*Rs can still depolarize the astrocyte to the same amplitudes as GABA*_A_*Rs and to larger amplitudes than Kir 4.1/GLT-1. Interestingly, NMDAR phase plots showed wider traces when compared to the GABA*_A_*R, mainly due to the large [K^+^]*_o_* independent depolarization (Fig 7 D vs. Fig 6 D). This contributed more to the efflux of K^+^ from the astrocyte (Fig C vs. Fig 6 C). The NMDAR model also matched quite well with the experimental results, suggesting that PAP depolarization amplitudes could be achieved without strong spillover or accumulation of neurotransmitters (Fig 7 E, Fig S10; SSE: 3.8). Our results indicate that increasing NMDAR/GABA*_A_*R had a strong impact on [K^+^]*_o_* and membrane potential, specifically seen as a wider loop structure in the phase plane. Our model suggests that NMDARs contribute most efficiently to depolarization, with the Kir 4.1 channel and NMDAR balance strongly determining the depolarization dynamics.

**Fig 7.**
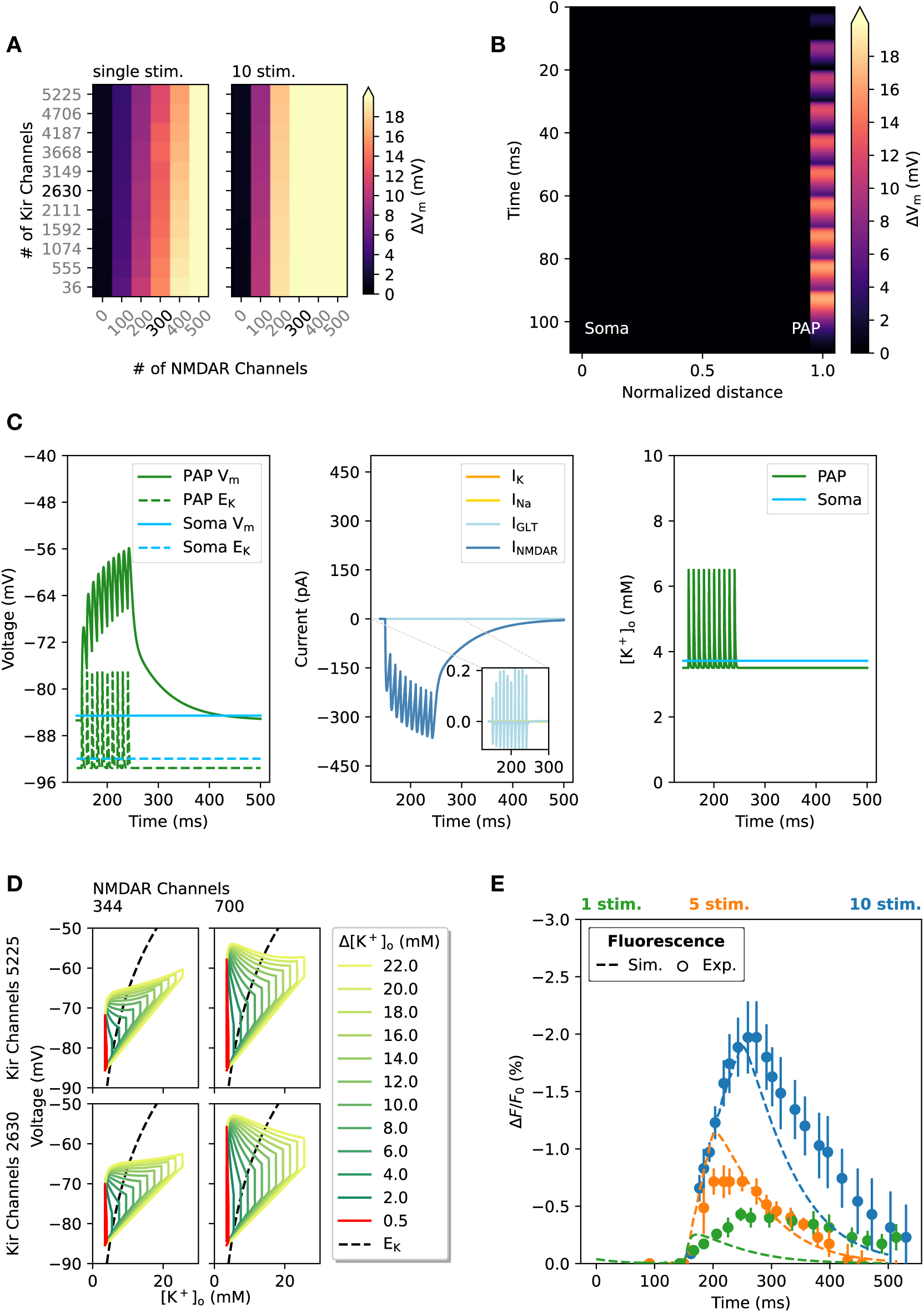
NMDAR driven depolarization **(A)** Heatmaps showing the typical depolarization response of PAPs with various counts of Kir 4.1 channels and NMDAR. The peak depolarization recorded at the PAP was compared for both Left) single and Right) 10 stimuli of 0.5 ms. All stimuli were followed by a 1 mM transient increase of glutamate. The colors indicate the membrane potential change (ГV*_m_*) for each condition, corresponding to the color scale within the figure. The black text labels indicate control conditions, while all gray values are models with modified channel numbers. **(B)** The voltage attenuation along the branch from soma to PAP over time, showing a complete lack of depolarization away from the PAP. Distance was normalized to the distance between the PAP far end and the soma. There are 344 NMDAR channels and 2630 Kir 4.1 channels to match control conditions used in panel A, C and E. **(C)** Left) Voltage (V*_m_*) along with reversal potential (*E*_K_) for PAP and soma, Middle) currents in the PAP (*I*_K_*, I*_Na_*, I*_NMDAR_*, I*_GLT_), Right) [K^+^]*_o_* within the PAP section and soma. Middle plot inset shows all currents other than *I*_NMDAR_ to show *I*_GLT_ has only a small contribution. **(D)** Phase planes of [*K*^+^]*_o_* vs. voltage for respective combination of Kir 4.1 and NMDAR counts. Traces for phases with different amplitudes of [K^+^]*_o_* stimuli are color-coded and labeled in the legend. Each panel contains four figures with the corresponding conditions labeled for each column and row. All phase planes show the model responses to a single stimulus. The control condition for the relative [K^+^]*_o_* increase that was used for typical sAPs is shown in red. The reversal potential, calculated from the voltage and [K^+^]*_o_*, is plotted as a broken black line. **(E)** Model fitting to experimental results observing peripheral depolarization in astrocytes [24]. The y-axis shows changes in the fluorescent voltage indicator (ArcLight;Г*F/F*_0_) for both simulation and experiment. Fitting was achieved by using channel counts as a free parameter (Fitted result: NMDAR 344). The fitting results are much better than with GABA inputs or the K^+^ spillover model (SSE: 3.8), with simulated peak amplitudes close to experimental ones for both 5 and 10 stimuli.

### Perisynaptic astrocyte processes respond to physiological single synapse activity

So far, our results suggest that potential mechanisms of local astrocyte depolarization can be classified into two large categories. The first category is Kir 4.1/GLT-mediated depolarization, which depends on high-frequency neuronal activity and the spillover of neurotransmitters and [K^+^]*_o_*. On the other hand, neurotransmitter-mediated depolarization shows amplitudes comparable to those of depolarization mediated by other mechanisms, without needing spillover. The differences between the two suggest that astrocyte depolarization responses could be heterogeneous depending on the channel composition of the PAP. We therefore examined how our model responds to synaptic input at near physiological activity, such as 50 Hz stimuli and theta burst stimulation (TBS). These protocols would be closer to physiological neuronal activity when compared to the 100 Hz sAP protocol. Using these various stimulation protocols, we tested four models explored in the previous passages, namely the K^+^ model, which only activated K^+^ channels, the GLT-1 model, which activated K^+^ channels and GLT-1, the NMDAR model, which activated both NMDAR and GLT-1 as well as K^+^ channels, and the GABA*_A_*R model, which activated K^+^channels and GABA*_A_*R. For a simple comparison, we uniformly activated 40 synapses, excluding soma and primary branch, to observe changes in both somatic membrane potential and the variance of synapses. As we examined responses to stimuli with a duration of 500 ms, which were much longer than in previous simulations, we considered the possibility of intracellular depletion for prolonged neurotransmitter-driven depolarization. Therefore, we included intracellular diffusion in these simulations. Furthermore, we also considered the slowing of diffusion dynamics in the synaptic cleft caused by a global increase in [K^+^]*_o_* and glutamate levels due to simultaneous activation, by implementing a decrease in time constants matching that of the experimental fitting for the spillover model (Fig 4 E).

As a result, we saw significantly different peak voltages among the stimulation protocols (one-way ANOVA; K^+^ model p-value: 0.99, GLT-1 model p-value: 0.99, GABA*_A_*R model p-value: 1.5×10*^→^*^2^, NMDAR model p-value: 4.9×10*^→^*^2^). At 50 Hz, neurotransmitter-mediated depolarization was smaller than at 100 Hz (Fig 8 A). Interestingly, this was not the case for TBS, suggesting that large local neurotransmitter-mediated depolarizations could occur with physiological inputs to the astrocyte (Fig 8 B). Specifically, for inhibitory synapses, we observed that amplitudes between 100 Hz and TBS were similar, predicting amplitudes observed with 100 Hz for inhibitory synapses are plausible with physiological stimuli. On the other hand, K^+^ and GLT-1 mediated depolarization did not change in peak amplitude regardless of protocol, suggesting that in tightly regulated synapses with minimal spillover, depolarization would not occur (Fig 8 C). We further observed that the isolation of these signals was maintained regardless of simultaneous PAP activation, which resulted in a *<* 1 mV change at the soma (Fig S12).

**Fig 8.**
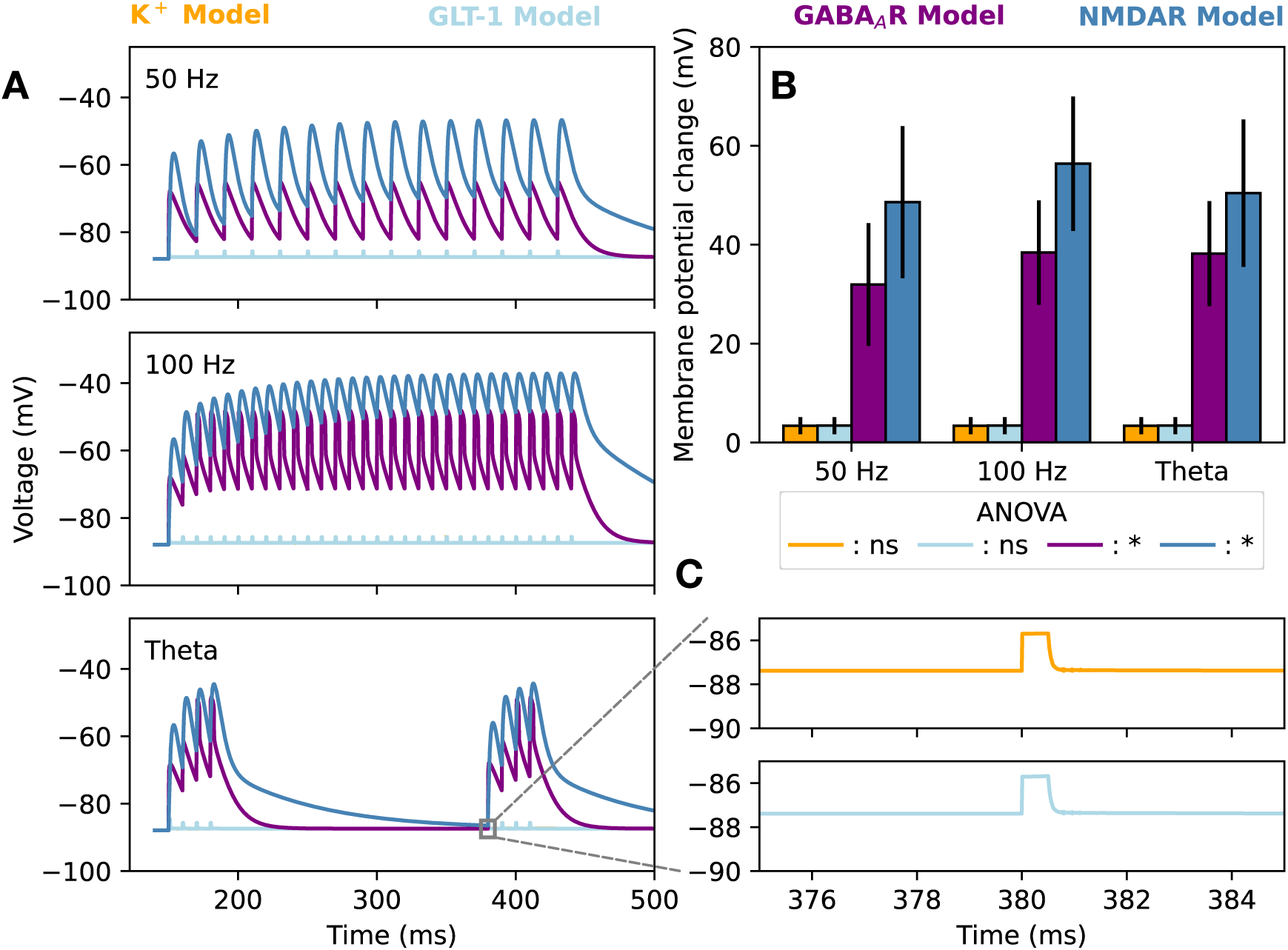
Physiological stimuli can depolarize astrocytes with neurotransmitters PAP depolarization induced by various stimulus protocols of 50 Hz, 100 Hz, and TBS for different models. We compared the K^+^ model (Fig 3), GLT-1 model (Fig 4), GABA*_A_*R model (Fig 6) and the NMDAR model (Fig 7). **(A)** Representative time course for each stimulus protocol showing differences in the accumulation of depolarization over time. **(B)** A bar plot showing peak membrane potential changes per model for each stimulus protocol (n=40). The NMDAR-mediated depolarization induced by the 100 Hz stimuli protocol had the largest mean peak depolarization amplitude. K^+^-mediated and GLT-1 mediated depolarization did not have any differences among stimulus protocols (ANOVA; K^+^ model p-value: 9.9 10*^-^*^1^, GLT-1 model p-value: 9.9 10*^→^*^1^, GABA*_A_*R model p-value: 1.5 10*^-^*^2^, NMDAR model p-value: 4.9×10*^-^*^2^). **(C)** Insets of single stimulus responses for the K^+^ model and GLT-1 model during TBS.

These results showcase key differences between the two types of astrocyte depolarization we observed in the model. Our results suggest that in case of physiological stimulation with minimal spillover, such as TBS, peripheral astrocytic depolarization is focal and induced only with neurotransmitters.

## Discussion

This study examined electrophysiological differences between the soma and periphery of cortical astrocytes using a new empirical, conductance-based mathematical model. Constructing a new conductance-based model with a focus on the peripheral compartment was necessary to implement realistic *in vivo* extracellular dynamics and discernible channel expression levels. This allowed for a detailed investigation of role of the Kir 4.1 channel, the major contributor to I_K_, and showed a clear distinction among somatic and peripheral electrophysiological properties.

We find that well-documented functions of astrocytes in K^+^ clearance and voltage attenuation are prominent at the whole cell level. Conversely, our model suggests that typical Nernstian properties do not apply for local stimuli, which are instead dictated by the local astrocyte subcompartment’s ability to reduce I_K_ escape. These properties highlight the importance of the axial conductance of local processes, suggesting that astrocyte depolarization amplitudes can be larger in tightly constricted portions of the cell, structures actually observed in astrocyte EM [58]. Our model also suggests that spatially clustered synapses can profoundly affect astrocytic depolarization amplitudes, enabling isolated astrocytic responses to local domains of activity. These mechanisms highlight a wide spectrum of potassium-mediated responses of astrocytes, indistinguishable by depolarization amplitude. Such potassium-driven responses observed in Armbruster et al. [24] are likely due to repetitive APs at the synaptic cradle or ECS pockets surrounding the axon, which we do not specifically distinguish in our model. Thus, further investigation into the relationship between the ECS and potassium diffusion in realistic morphologies is expected to yield deeper insight into local astrocyte depolarization and neuronal excitability. Throughout our results, our model shows the sponge-like capabilities of astrocytes to handle local K^+^ at the soma without large changes in membrane potential, which matches experimentally observed properties [22].

Although we found Kir 4.1 to be the major driver for K^+^-mediated depolarization, consistent with previous literature [22, 33, 59], a typical isolated PAP depolarization of 20 mV required *→* 22 mM of K^+^ for physiological Kir 4.1 densities. This concentration is typically considered a large [K^+^]*_o_* change, which exceeds the predicted physiological upper bound of [K^+^]*_o_* changes [60–62]. We believe this is true regardless of small ECS as well, with our results showing the difficulty of achieving large responses with single presynaptic events (Fig 3 B).

We further examined this model with neurotransmitters to examine how K^+^-independent mechanisms would respond differently between somatic and peripheral regimes. Our examination of GLT-1 indicated that they contribute only in whole-cell configurations, unlike the other ionotropic receptors we examined. Furthermore, we saw that significant K^+^/GLT-1 depolarization required strong accumulation and spillover of neurotransmitters (Fig 4). These results conform with experimental evidence of astrocyte depolarization during epilepsy [63]. Alternatively, our model also suggested depolarization mechanisms independent of spillover, mediated by ionotropic neurotransmitters. These mechanisms show different ways to obtain isolated depolarization, which may indicate a more active role of astrocyte depolarization, tailored to individual synapses.

We believe that our model highlights two novel insights into astrocytic electrophysiology. First, dynamics in whole cells and the local periphery are vastly different, with conventional notions like the Nernst potential being insufficient at the nanoscopic level (Fig 1 D vs. Fig 2 C). We also suggest that local electrophysiological properties are governed by local conductance ratios that determine the astrocyte membrane’s ability to prevent escape of I_K_. Second, astrocytes may use ionotropic synaptic receptors *in vivo* to alter membrane potentials, thereby significantly coupling astrocyte responses to neuronal stimuli (Fig 6, 7). Although many *ex vivo* protocols suggest the contrary [64, 65], our model indicates that astrocyte conductance and morphology may strongly mask synaptic receptor currents in somatic voltage clamp studies (Fig 5). These insights together imply that astrocytes may have dynamic neuronal responses that are invisible to conventional electrophysiological techniques. We believe that, although the study by Armbruster et al. [24] was the first step toward identifying these hidden astrocyte responses, a more thorough examination that distinguishes whole-cell and local differences is necessary. Our model indicates that K^+^ mechanisms alone in the periphery require high-frequency stimulation and strong ion and signaling molecule accumulation in the ECS, which may not be a common phenomenon in physiological brain conditions. Indeed, in vivo recordings of mice somatosensory cortex L2/3 during running show ≈ 1 mM increase in [K^+^]*_o_*, highlighting the difficulty to achieve confined large amplitudes of K^+^ in physiological settings [66]. Notice also that results from Armbruster et al. [24] omit the possibilities of astrocytic synaptic receptors because they apply APV and DNQX for all experiments and only stimulate excitatory axons. Their experiments to measure specific channel contributions were analyzed using whole image averages, neglecting possible distribution differences within nanoscopic domains. Any receptor that would be clustered in peripheral subcompartments of the astrocyte would be neglected, as the membrane area is much smaller than that of the whole cell. This leaves the possibility for further examination focused on local astrocytic responses to neuronal activity, using techniques with better spatial resolution, such as expansion microscopy or stimulated emission depletion (STED) imaging. The apparent absence of effect of APV on the experimentally observed ROI depolarizations [24] may also be due to differences in astrocyte NMDAR subunit composition, as certain NMDAR subunits are less susceptible to APV [67], which may also account for reports on astrocyte NMDA-induced, APV-insensitive currents [68, 69]. The low and localized expression of astrocytic NMDAR has been discussed previously [70, 71].

We also re-examined the limitations of fluorescent voltage indicators in our model using simulated fluorescence. Our simulated fluorescence reflected experimentally observed responses to various voltage-clamp durations, highlighting the limitations of fluorescent voltage indicators such as ArcLight for fast transient responses (Fig S9A). The response of fluorescent voltage indicators was heavily influenced by the time dynamics of voltage responses, clearly reflected in our experimental comparisons (Fig 3 D, Fig 4 E, Fig 6 E, Fig 7 E, Fig S9 B,C). One example of such signal dynamics affecting fluorescence readout is the experimentally observed buildup of fluorescence observed for AP trains [40]. This aspect is clearly reflected in our simulation results of the single synapse model (Fig 3D). Interestingly, our results indicate that the apparent fluorescent buildup due to voltage indicators alone is insufficient to account for the depolarizations observed in Armbruster et al. [24], suggesting the presence of a slower depolarization component independent of the fluorescent artifacts. In this regard, our model suggests that both slow neurotransmitter activation and spillover can produce these effects (Fig 4 E, Fig 7 E).

Our model, like many others, neglects the effects of glial swelling induced by neuronal activity. In experimental studies, K^+^ influx through the glial membrane strongly contributes to glial swelling, presumably due to the coupling between Kir 4.1 and aquaporins (AQP4) [72]. In theory, the ECS volume would decrease due to local glial swelling, which could alter [K^+^]*_o_*. Current technical limitations, as well as limited experimental evidence on K^+^ dynamics in the nanoscopic regime, inhibit a thorough investigation of the impact of glial swelling on synaptic efficacy in our multicompartment model. Although multi-domain models with simplified geometry exist, these studies simulate only osmotic pressure gradients, which constitute only a limited component of glial swelling. The development of a dynamic volume-morphing model of glial processes would be interesting for further investigating K^+^ dynamics in response to neuronal activity.

PAP depolarization may also strongly alter astrocytic function in information processing [24]. This could be achieved by the voltage dependence of GLT-1, as well as contributing to calcium dynamics by altering Na^+^/Ca^2+^ exchanger functionality [73]. However, the extent to which these mechanisms affect brain computation is unknown. The prevailing assumption is that astrocyte computation is done through increases of intracellular calcium [74, 75]. Although, at first glance, PAP depolarization may seem to drive calcium influx, we believe depolarization does not directly activate voltage-dependent calcium channels (VDCC). This is because major VDCCs on astrocytes (L-type, N-type, R-type) have high voltage thresholds that are difficult to achieve with ranges of depolarization shown experimentally [24] and in the model [76]. Previous experimental results match this description, where astrocyte L-type VDCC-mediated influx required bath application of 75 mM [K^+^]*_o_*, expected to cause large membrane depolarization [1].

Recent studies also reveal the capabilities of astrocyte-mediated territorial K^+^ control at the single-cell level [77]. Our model suggests that this function could also exist at the synapse level, as indicated by astrocytes’ ability to isolate and respond to neuronal input. Our model predicts local astrocytic K^+^ efflux in response to neuronal activity, contributing to neuronal excitability at the synaptic level. This opens the door to broader possibilities for how K^+^ regulation may contribute to synaptic function, along with other well-established astrocyte contributions, such as gliotransmission and metabolic signaling [78–81]. If so, local protein expressions in astrocyte subcompartments would be more crucial in understanding astrocyte functions, in comparison to global expression profiles. Given this possibility of dynamic astrocyte synaptic responses, we believe it would be possible for the astrocyte to tailor its response based on the specific synapse it is in contact with. For example, the number of NMDARs in each PAP may be actively regulated by protein trafficking, where specific PAPs are tailored to have more NMDARs. Protein trafficking in astrocytes is not well understood, but certain proteins like glutamate transporters and AQP4 have been observed to be trafficked via anterograde vesicle transport [82, 83].

In summary, our results suggest vastly different electrophysiological responses dependent on spatial scale. Our model predicts the breakdown of Nernstian properties of astrocytes for local responses driven by synaptic activity. Our current findings suggest that PAP depolarization can be triggered by both high-frequency stimulation and synaptic receptors. With the whole-cell conductance-based model, we showed K^+^/GLT-1 mediated mechanisms required strong spillover and high-frequency stimulation, while synaptic receptor-mediated depolarization could occur with spatial isolation. For K^+^/GLT-1 mediated depolarization, we show that depolarization requires K^+^ amplitudes that are much higher than initially anticipated by previous reports [24]. As the impact of PAP depolarization on synaptic plasticity is still unknown, further investigation into the properties of K^+^ and glutamate diffusion at the nanoscopic spatial scale is required.

### Model design

The objective of the study was to construct a realistic whole-cell astrocyte *in silico* model, capable of recreating electrophysiological responses observed in both somatic and peripheral regimes. Astrocyte depolarization is strongly dependent on [K^+^]*_o_*, which requires the model to dynamically update the Nernst reversal potential. This excluded the use of channel models that included built-in mechanisms for maintaining membrane potential, which are often used to model astrocyte electrophysiology [33]. The morphology used in this model was from Savtchenko et al., which used hippocampal CA1 protoplasmic astrocytes [29]. Morphological differences between protoplasmic hippocampal astrocytes and cortical somatosensory astrocytes are minimal, with high correlation of morphological characteristics between the two [84]. Intracellular diffusion mechanisms did not affect the results of localized short pulse trains (Fig S5), and therefore were clamped for most simulations. Other simulations with longer pulse trains were simulated with intracellular diffusion. The model morphology was defined in NEURON [85], in the file GeometryAstrocyteCA1.hoc. See also Fig S1 A,B for additional morphological details. All channel models were defined in NEURON MODL [85]. All parameters used in the model are listed in Table 1 and Table S1-S11. Any alteration of channel counts was achieved by linearly varying the single-channel conductance. For pumps in the model, such as the Na/K-ATPase (NKA) we altered the current density instead. Kir 4.1, glutamate transporters (GLT-1), K^+^leak currents (TREK-1/TWIK-1), and sodium leak channels were uniformly distributed for all simulations based on the channel counts defined in the PAP. For GABA*_A_*R and NMDAR, any change was directly reflected in channel numbers solely in the PAP. Kir 4.1 and leak K^+^ channels (TREK-1/TWIK-1) were fitted to experimental data shown in Fig S2. All neurotransmitter-related channels, including GLT-1, were implemented as point processes to recreate neurotransmitter release, with synaptic weight converted to concentration, used for Hill-dependent activation. NKA, TREK-1/TWIK-1, and sodium leaks were defined as membrane mechanisms. All referenced MODL files can be found in the neuronMOD file within the uploaded repository.

**Table 1.** Constants for whole-cell astrocyte model.

| Parameter | Value | Units | Description | Comment/Ref. |
| --- | --- | --- | --- | --- |
| $R_a$ | 100 | $\Omega \text{ cm}$ | Axial resistance | [29] |
| $C_m$ | 0.8 | $\mu\text{F cm}^{-2}$ | Membrane capacitance | [29] |
| $g_L$ | $6.89 \times 10^{-4}$ | $\text{S cm}^{-2}$ | Passive conductance | [29] |
| $E_L$ | -85 | mV | Reversal potential of passive leak current | [29] |
| $T$ | 307 | K | Temperature matching experimental setup | [24] |
| $R$ | 8.3145 | $\text{J K}^{-1}$ | Gas constant | |
| $F$ | 96485.3321 | $\text{C mol}^{-1}$ | Faraday constant | |
| $[\text{K}^+]_i$ | 120 | mM | Intracellular potassium concentration set for potassium reversal potential of -91.7 mV | [3] |
| $[\text{K}^+]_o^b$ | 3.5 | mM | Extracellular potassium concentration baseline | [24] |
| $[\text{Na}^+]_i$ | 15 | mM | Intracellular sodium concentration | [100] |
| $[\text{Na}^+]_o$ | 140 | mM | Extracellular sodium concentration | [100] |
| $[\text{Cl}^-]_i$ | 30 | mM | Intracellular chloride concentration | [100] |
| $[\text{Cl}^-]_o$ | 130 | mM | Extracellular chloride concentration | [96] |
| $[\text{Mg}^{2+}]_o$ | 1 | mM | Extracellular magnesium concentration | [101] |
| $[\text{Glu}^+]_i$ | 0.3 | mM | Intracellular glutamate concentration | [29] |
| $[\text{Glu}^+]_o$ | $20 \times 10^{-6}$ | mM | Extracellular glutamate concentration | [29] |
| $g_{\text{K},L}$ | 1.15 | $\text{S cm}^{-2}$ | Leak potassium conductance fit to experimentally measured space constants | [22] |
| $l_{\text{ecs}}$ | 40 | nm | Depth of ECS compartment | [102] |

### Extracellular space

The ECS was modeled as a shell around the cellular compartment with an effective thickness of 400 Å at the PAP, and 1000 ^°^A at the soma. These values roughly cover the ranges observed in experimental measurements and were linearly interpolated based on compartment diameter, in the aforementioned range [86, 87]. The extracellular diffusion between sections was not considered, as the selected PAP regions were thought to be isolated. The time constant for flux out of the extracellular compartment was calculated from diffusion, based on Fick’s law (See SI text S1)

### Intracellular Diffusion

Intracellular diffusion was modeled using the framework from Savtchenko et al. [29]. Given the high [K^+^]*_i_* and its relatively rapid diffusion, no radial shells were implemented; only longitudinal diffusion was considered. Longitudinal diffusion among compartments was implemented following chapter 9 of the NEURON book [85]. Experiments with dynamic [K^+^]*_i_*showed significant K^+^ depletion within the cell, especially in compartments with large surface areas. Therefore, we specifically refit NKA maximum amplitudes to retain RMP at -86 mV.

### Ion current to concentration conversion

The K^+^ flux from intracellular to extracellular compartments, and vice-versa was calculated with the conversion,

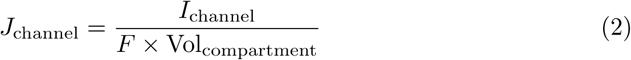

where *J, I* describe the flux and current of any channel, *F* is the Faraday constant and Vol is the volume of the compartment. For K^+^, this was modeled as a NEURON MODL file that follows chapter 9 of the NEURON book [85], modeling extracellular Ca^2+^ concentrations and replacing Ca^2+^ with its respective ion properties. The ion flux conversion files were defined as potassiumAccum.mod. Chloride and sodium concentrations in the extracellular/intracellular space were constant, and we did not consider local changes.

### Kir 4.1

The Kir 4.1 channel model was adapted from Yim et al. [88] with a [K^+^]*_o_*dependent square-root law [89]. This model was chosen over previous models [33], which included arbitrary RMP sinks that could skew I*_K_* amplitudes.

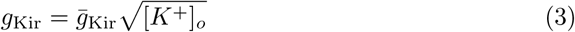

where *g̅*_Kir_ is the maximum conductance. Maximum conductance was calculated to fit single-channel recordings and [K^+^]*_o_* of that experiment [90]. Details can be found in the Kir2.mod file and SI text S2.

Estimates of Kir 4.1 channel densities, along with passive leak conductances, were derived to match experimental values that predicted space constants at the soma [34]. Specific derivations are outlined in SI text S3 and Fig S3.

### K2P leak channels

The K2P TREK-1 and TWIK-1 leak channels were implemented as a typical passive K^+^ leak channel to reflect both channel properties. Details can be found in the SI text S4.

### Na/K-ATPase

The Na/K-ATPase (NKA) model was based on the work of Somjen et al., which utilized an analytical Hill-type reaction scheme for ion pumping [26]. The maximal current was implemented to match previous model parameters of 1 mA cm*^→^*^2^ [26]. Details can be found in the SI text S5.

### NMDAR

The NMDAR model is adapted from Moradi et al. [91] with a few alterations to fit astrocyte NMDAR recordings. First, the voltage dependence of time constants for the NMDAR model was refit to that of astrocyte NMDAR recordings by [12]. Second, the magnesium dependence of the model was reduced, as astrocyte NMDAR currents are independent of extracellular magnesium concentrations [12]. Lastly, the maximum single-channel conductance was calculated from Jahr et al. [92]. This was under the assumption that NMDAR resides primarily at the fine processes [93]. PAPs during spillover experiments contained the most NMDAR at the initial 0.3 µm length and declined with units of 0.3 µm. Details can be found in the SynExp5NMDA.mod file and SI text S6.

### GABA*_A_*R

The GABA*_A_*R model was a classic two-state synaptic conductance model adapted from Schulz et al. [94], with the reversal potential adjusted to fit astrocytic conditions. As GABA*_A_*R reversal potentials were that of chloride [95, 96], we calculated the chloride reversal potential from intra/extracellular concentrations using the Nernst equation. This resulted in a reversal potential of approximately -40 mV. Details can be found in the inh.mod file and SI text S7.

### Glutamate Transporter

The glutamate transporter model (for GLT-1) utilized in this study was modified from that used in Savtchenko et al. [29] and converted the original membrane mechanism to a point process. Furthermore, the model was specifically corrected to affect K^+^ dynamics and the I_K_, as the channel was observed to have [K^+^]*_o_* and voltage-dependent uptake [97]. The resulting model showed the expected current amplitudes observed in experiments (Fig 5 C). Details can be found in the GluTrans.mod file and SI text S8.

### Leak channels

Leak channels for Na^+^ were modeled as a generic passive channel with an analytically calculated conductance. The specific conductance value was analytically calculated so the RMP was -85 mV. Further explanations of the calculations are detailed in SI text S9. The ion leak mechanism was defined as naleak.mod.

For ion independent passive membrane properties, the leak was modeled as passive currents to match experimentally observed conductances [29].

### Gap junctions

Gap junctions in the model were implemented as passive shunts, reflecting measured experimental trans-junctional conductances of the astrocyte. Assuming tiling of astrocytes and minimal overlap between adjacent astrocytes, all gap junctions were positioned only at the peripheral end compartments of the model. Inclusion of gap junctions controlled astrocyte potassium clearance in bath application protocols, matching general experimentally observed characteristics (Fig S4). Further explanations of conductance values can be found in SI text S10.

### Calculating local current escape

The depolarization in local compartments was strongly correlated with the parameter Г*_K_*defined as follows,

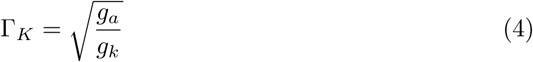

where *g_a_* was the axial conductance per cross-sectional area, and *g_k_* was the specific K^+^ membrane conductance. Each of these conductances was calculated from two experimental protocols: measurements of input resistance and measurements of minimum [K^+^]*_o_* change required for fixed amplitude current generation. Using each minimum required Δ[K^+^] (relative to baseline [K^+^]*^b^*) to induce 0.05 nA (Δ*I_K_*) we calculated total K^+^ conductance (*G_k_*) as,

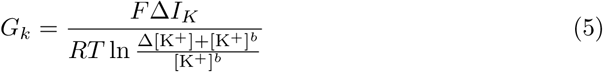

where *R, T, F* are gas constant, temperature, and Faraday constant, respectively. Utilizing this value and local input resistance (*R_i_*), assuming membrane conductances are dominated by K^+^, the total axial conductance (*G_a_*) was calculated using Kirchhoff’s law as,

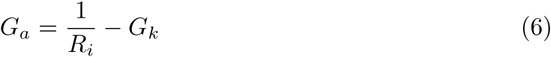

We then converted these total conductances to specific K^+^ membrane conductance and axial conductance per cross-sectional area by,

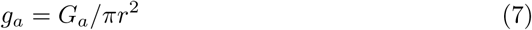

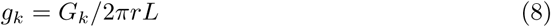

where *r, L* are the radius of the compartment and the length of the compartment. These values were used to compute Г*_K_*.

### Stimulation Protocols

Before stimulation, the model underwent an equilibrium phase during which [K^+^]*_o_* and its associated currents reached a steady state. This equilibrium phase lasted 150 ms, during which all channel conditions reached equilibrium. Bath application and *in vivo* stimulation protocols were recreated in the model by clamping [K^+^]*_o_* for appropriate durations. Changes in K^+^ for *in vivo* stimulation followed experimental results found in Chever et al. [2]. Either localized sections throughout the cell or the entire cell were targeted for local and global stimulation, respectively. Plots of single PAPs in this paper used a seed value of 1 to randomly select a location in the astrocyte morphology, unless stated otherwise. This seed was chosen as it was a PAP with the median response to K^+^ stimuli (Fig 2, Fig S8).

Stimulation protocols using simplified action potentials (sAPs) to depolarize astrocytes were performed by altering conductances in synaptic receptor models and changing [K^+^]*_o_*. The sAP stimuli were triggered at PAP-designated astrocyte sections at 100 Hz to match experimental protocols [24], unless stated otherwise. Each stimulation was broken down as a 0.5 mM [K^+^]*_o_* increase that lasts for 0.5 ms and a 1 mM glutamate or 1.5 mM GABA increase unless stated otherwise. Glutamate and GABA-induced changes in NMDAR and GABA*_A_*R were implemented as Dirac delta-function changes in neurotransmitter concentrations, with neurotransmitter diffusion dynamics integrated into the model itself, as is typical for NEURON-implemented synaptic conductances. For GLT-1, glutamate concentrations followed a double exponential time-course of synaptic glutamate rise-decay dynamics, defined by Savtchenko et al. [29] (See Table S9). The K^+^ increases are confined to the ECS, at locations defined as the PAP.

### Model fitting to experimental results

K^+^ conductance for Kir 4.1 and passive K^+^channels were fit to experimental results [22, 34]. For ionotropic receptors, the model was fit to experimental results in Armbruster et al. [24] by selecting unconstrained parameters such as channel numbers, K^+^ amplitude, and PAP size. Explicitly for the GLT-mediated depolarization model with spillover, the glutamate decay time constant was additionally considered to be a free parameter. Fitting was performed using the Nelder-Mead gradient search algorithm in the Scipy package [98] with arbitrary initial values. For all other channels, conductance, channel density, and kinetics were constrained by previous experimental results.

### Simulations and analyses

All models were implemented in NEURON, where the default astrocyte model was constructed using NEURON HOC version 8.2.0. Specific simulation programming was done by accessing the HOC object in the Python interface using custom code. All ODEs were solved using NEURON’s variable integrating step (CVODE) solver, unless stated otherwise. MPICH [99] was used to parallelize individual simulations, using a custom code for repetitive simulations.

## Acknowledgements

We thank the lab members for their helpful insights on our manuscript.

