## Supplemental Figure, Table, Information for "Global Nernstian astrocytic depolarization breaks down during local synaptic input"

**This PDF file includes:**

- Fig S1 to S12
- SI text S1 to S12
- Table S1 to S11

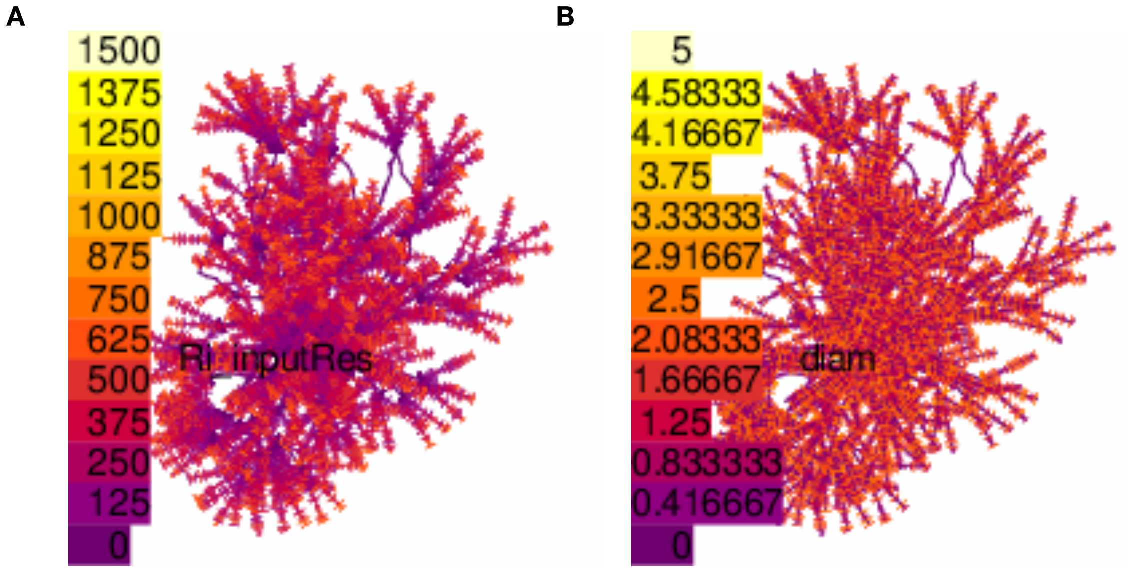


**Fig S1.** Morphological characteristics of the astrocyte whole-cell model
**(A)** Input resistance (color scale 0∼1500 MΩ) **(B)** diameter of each section (color scale 0∼5 µm) of the hippocampal astrocyte morphology. Morphological differences between protoplasmic hippocampal astrocytes and cortical somatosensory astrocytes are minimal, with high correlation of morphological characteristics between the two [84].

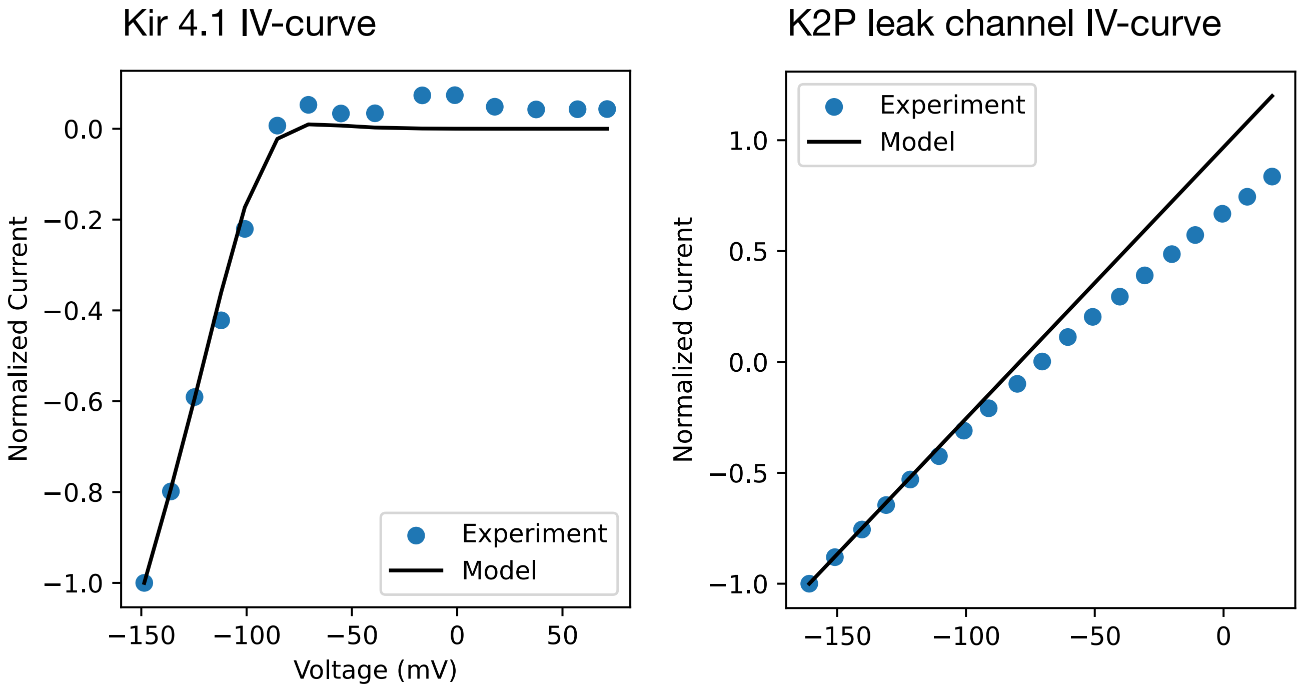


**Fig S2.** Comparison of experimental data to channel models
Plots showing the normalized current-voltage relationships for Left) Kir 4.1 and Right) TREK-1/TWIK-1 passive leak currents. Each channel was fit to experimental results, where Kir 4.1 fits to Kir 4.1 channel experimental data [103, 104] and TREK-1/TWIK-1 fits to experimental data used in Zhou et al. [3]. The model fits the data well for the voltage ranges used in this study. All experimental conditions, such as intra/extracellular $K^+$ concentrations and temperature, are considered in the model and were matched with the experimental data.

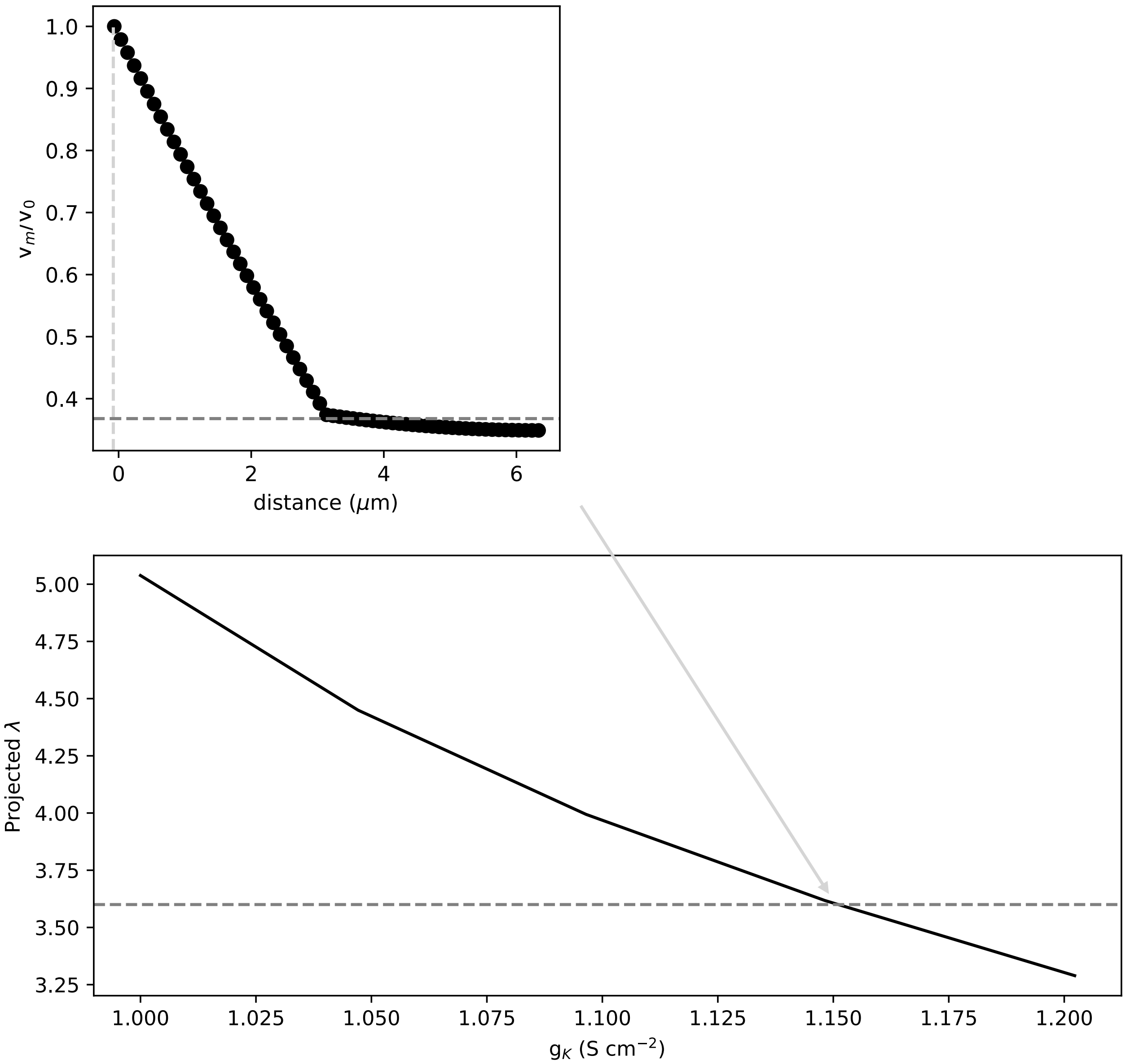


**Fig S3.** Model fitting to dual patch experimental results

Determining the total $K^+$ conductance to recreate experimentally predicted space constants measured at the soma using dual patch-clamp techniques. The model also includes sodium conductances, calculated to maintain RMP. Top) The voltage attenuation trace for $g_K = 1.15\,\mathrm{S\,cm^{-2}}$. We measured voltage attenuation ($V_m/V_0$) when the voltage was clamped at one end of the soma and fit the model to match the $1/e$ (dark gray horizontal dotted line) reduction from the holding potential. The location of the patch clamp was defined as 0, and the closer edge of the soma is indicated by the light gray vertical dotted line. The sudden change in attenuation at about 3 μm is due to the location of attached primary branches within the morphology. Bottom) The projected space constant ($\lambda$) for various $K^+$ conductance values in the model. The broken line represents the experimentally measured space constant of 3.6 μm.

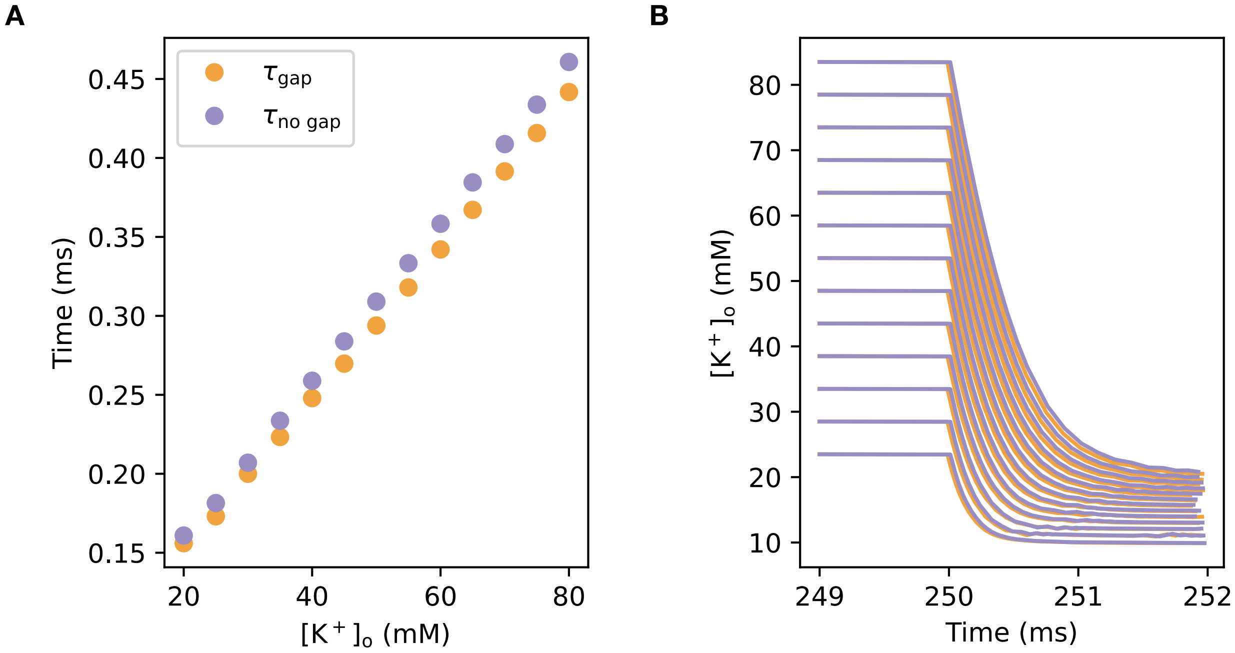


**Fig S4.** Effects of gap junction inhibition on $K^+$ clearance during bath application
The measured time constants ($\tau$) of $[K^+]_o$ clearance during bath application of various amplitudes. $[K^+]_o$ was clamped for 10 ms after initialization. The model included intracellular diffusion, $K^+$ channels, and gap junctions. Extracellular diffusional leak was turned off during the simulation to reflect bath application protocols. **(A)** $\tau$ for $K^+$ clearance for respective models. The model with gap junctions (orange) has consistently shorter $\tau$ for $K^+$ clearance than the model without gap junctions (purple). Additionally, the difference in $\tau$ increases when $[K^+]_o$ amplitudes are larger. **(B)** Traces of $K^+$ clearance for each respective stimulus, superimposed. All traces from the no gap junction model (purple) are slightly slower than their respective counterparts (orange), which is more prominent for larger $K^+$ amplitudes.

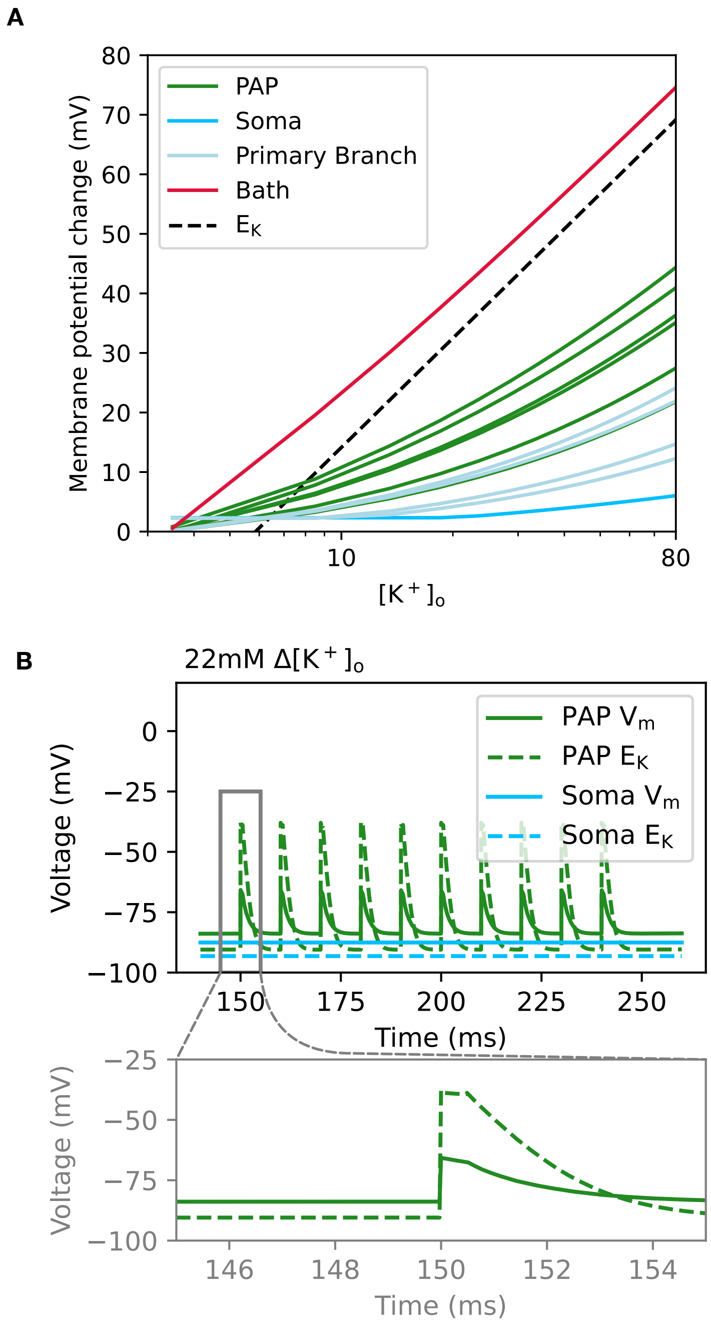


**Fig S5.** PAP depolarization in model with intracellular diffusion

**Fig S5.** PAP depolarization in model with intracellular diffusion
**(A)** Depolarization response to $K^+$ changes for soma, PAPs, and primary branch with intracellular diffusion turned on in the model. Most of the responses align with the results of the no intracellular diffusion model (Fig. 2C), except for a slightly stronger coupling to reversal potential for bath application (in $[K^+]_o$ stimuli over 10 mM), and very small increases of localized responses for primary branch and soma. Sampling of PAPs was reduced to decrease computational expense. **(B)** Astrocyte membrane potential response to 22 mM $[K^+]_o$ stimulus, with intracellular diffusion. The effects of intracellular diffusion are minimal, as the intracellular volume is larger than that of the extracellular volume, and $[K^+]_i >> [K^+]_o$.

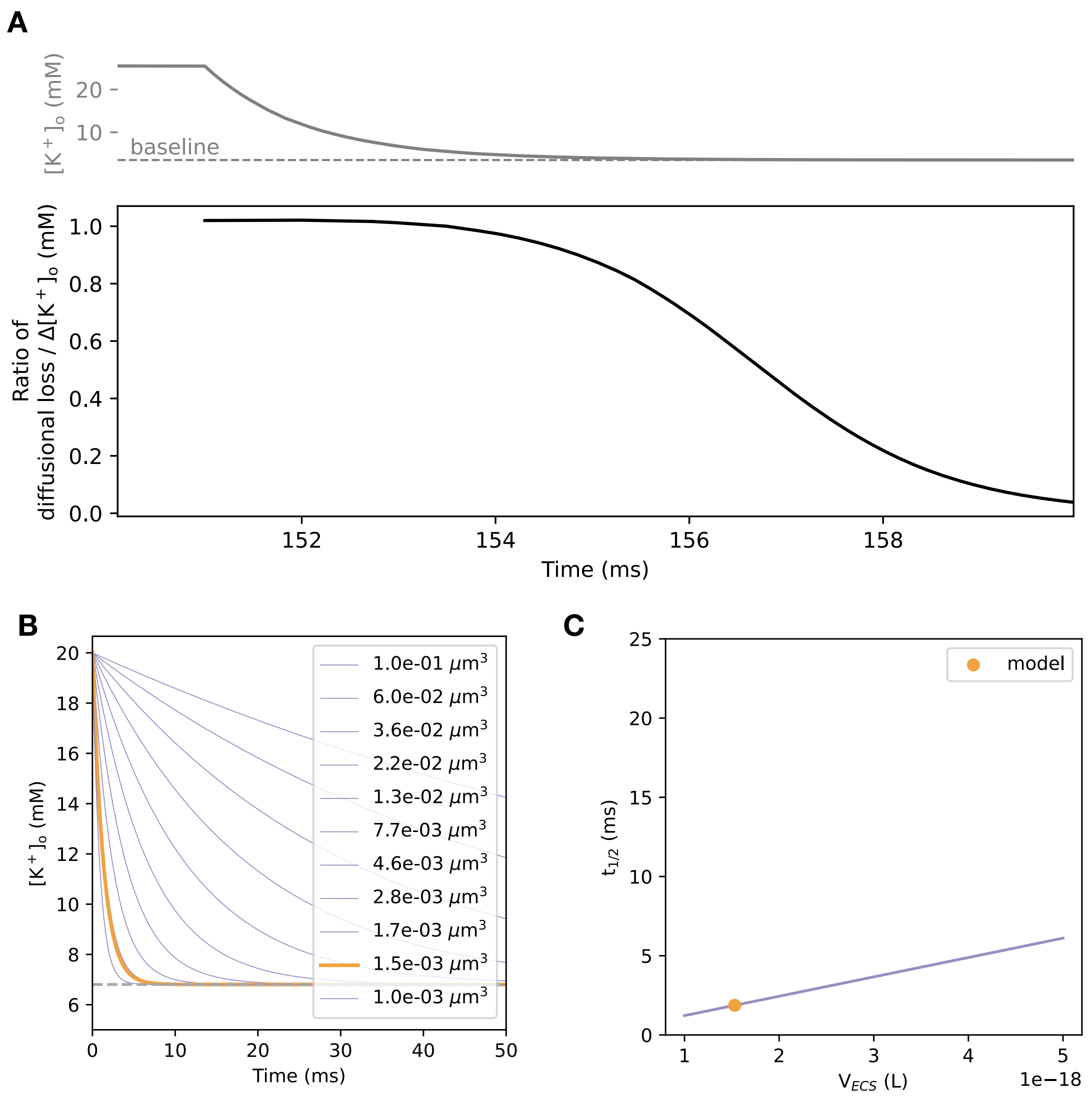


**Fig S6.** $K^+$ clearance in the synaptic cleft
**(A)** Ratio of $K^+$ clearance from the extracellular space (ECS). Top) The total $[K^+]_o$ (gray). Bottom) The ratio between extracellular diffusional loss from the ECS and the total change in $[K^+]_o$ (black). The ratio plot shows the astrocyte response to 22 mM $\Delta[K^+]_o$ with no glutamate. $K^+$ is increased at 150 ms for 1 ms and the ratio is recorded afterward. Extracellular diffusion dominates the $[K^+]_o$ clearance until $[K^+]_o$ relaxes to baseline. **(B)** Results for analytical model for Kir 4.1 mediated uptake of $[K^+]_o$. Model derivation is detailed in SI text S11. Results indicate that shrinking the ECS promotes faster dynamics of $K^+$ change in the extracellular compartment. The trace for ECS sizes comparable to our model is plotted as an orange line. All other ECS sizes are plotted in purple. The gray dotted line expresses the $[K^+]_o$ at equilibrium for the analytical model. **(C)** Half-life for $K^+$ decay plotted for various ECS sizes using the analytical model. The ECS space used in our conductance-based model is indicated as an orange circle within the plot. All other conditions are plotted in purple. These plots indicate a short half-life of extracellular $K^+$ mediated purely by Kir 4.1 uptake in diffusion-inhibited conditions.

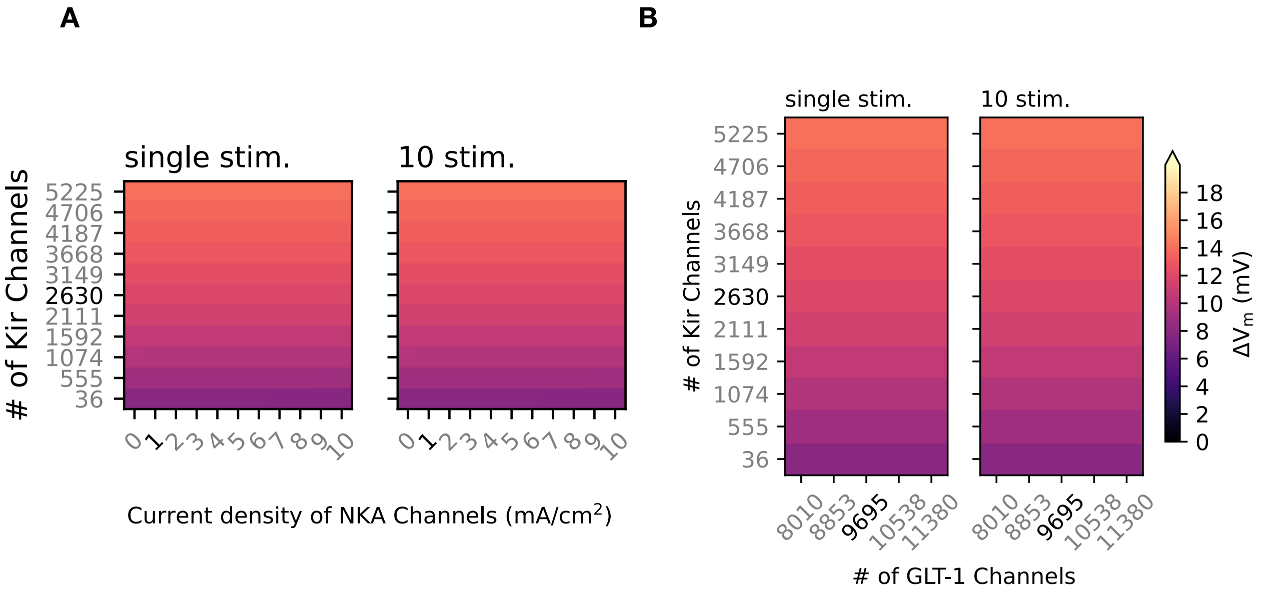


**Fig S7.** Effects of changing NKA or GLT-1 channel densities are minimal compared to Kir 4.1 changes Compares the contributions of Left) Na/K-ATPase (NKA) and Right) glutamate transporters (GLT-1) to $K^+$ uptake and membrane potential depolarization. **(A,B)** Heatmaps showing maximum depolarization in response to left) 1 right) 10 pulses of sAP with amplitude of 10 mM $K^+$ stimuli for changes in densities of A) NKA or B) GLT-1. Heatmaps compare differences in counts of Kir 4.1 channels and the other astrocyte channels. The heatmap shows peak depolarization ($\Delta V_m$) recorded at the PAP. The colors indicate the membrane potential for each condition, corresponding to the color scale within the figure. The black text label indicates the control conditions (Kir 4.1: 2630 channels; NKA: 1 mA cm$^{-2}$ [26]; GLT-1: 9695 channels in the PAP [46]), and gray colors are adjusted values. Both plots indicate strong dependence on Kir 4.1 densities rather than those of NKA or GLT-1 for transient potassium bursts (sAP).

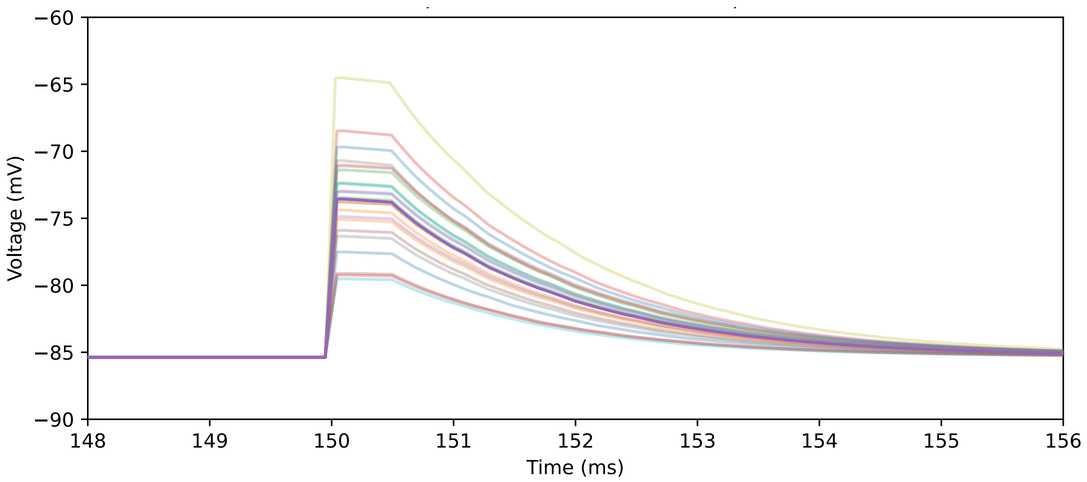


**Fig S8.** Variability of PAP responses dependent on location

Variability of PAP responses to local $[K^+]_o$ changes is dependent on location. Each location was selected at random with different seeds. Individual plots for PAP voltage response to 10 mM $[K^+]_o$ changes. A depolarization of 20 mV was achieved in only one PAP, which had the lowest $\Gamma_K$ (Fig 2 E). Each trace corresponds to a different seed, with seed 1 emphasized.

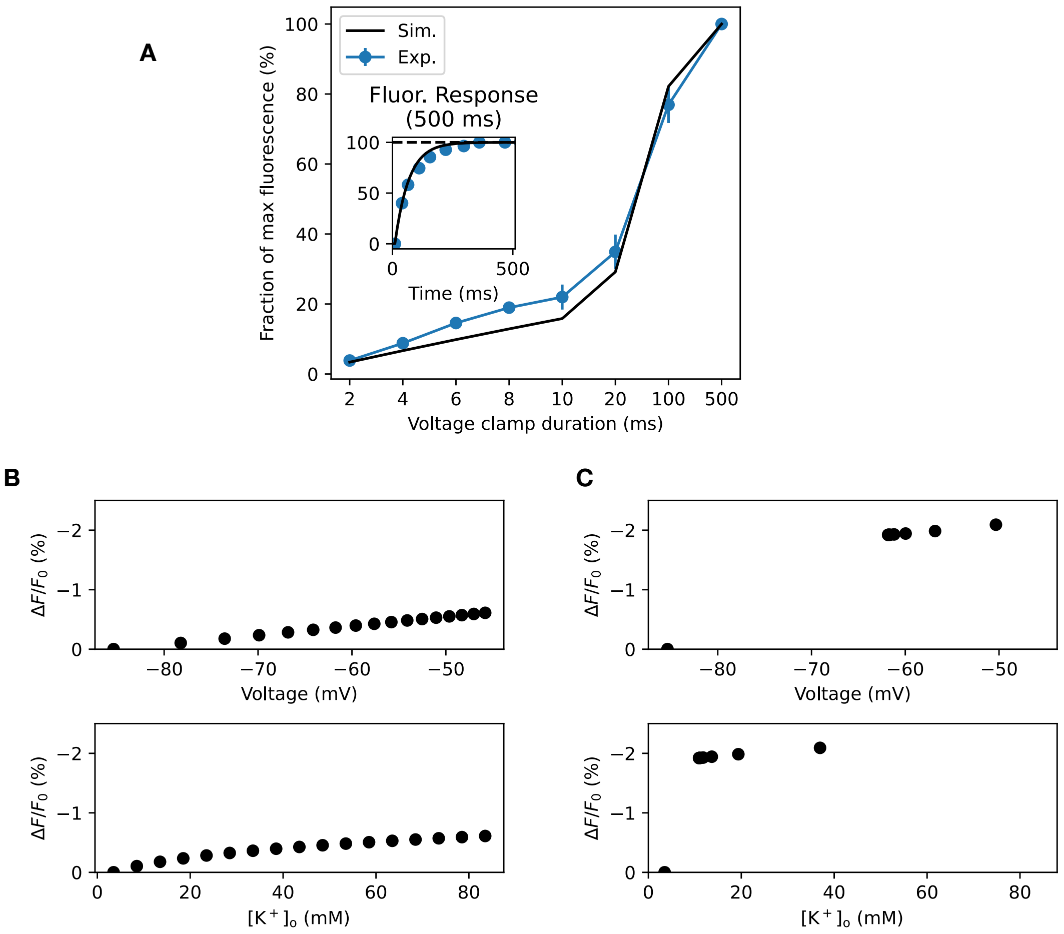


**Fig S9.** Simulation of fluorescence signals
**(A)** Comparison of simulation to experimental *in vitro* response of GEVI ArcLight, during voltage clamp of 100 mV with various durations [40]. The simulated fluorescence was fit to the time dynamics of the fluorescence response to 100 mV voltage clamp for 500 ms (inset figure). The experiment evaluates the fraction of the maximal response the GEVI ArcLight can indicate for each voltage clamp duration, with 500 ms necessary to fully reflect the 100 mV response. The simulated responses roughly follow the results of the experiment (2,4,6,8,10,20,100,500 ms). These results show the slow activation of the GEVI ArcLight, which will filter our transient responses with durations below 10 ms.
**(B,C)** Simulated fluorescence responses for various $K^+$ stimuli (bottom) with conditions matching the experimental fit B) Single synapse model (Fig 3 D), C) Spillover model (Fig 4 E). For the single synapse model, the response of peak voltage to fluorescence is linear (top), although the calibration slope is much smaller than the one estimated by Armbruster et al. [24]. On the other hand, the $K^+$ response under the spillover context in much more non-linear, with a sudden jump of response for $K^+$ change larger than 0.01 mM. The slope, after the jump, is much smaller than the linear calibration curve observed in Armbruster et al. [24].

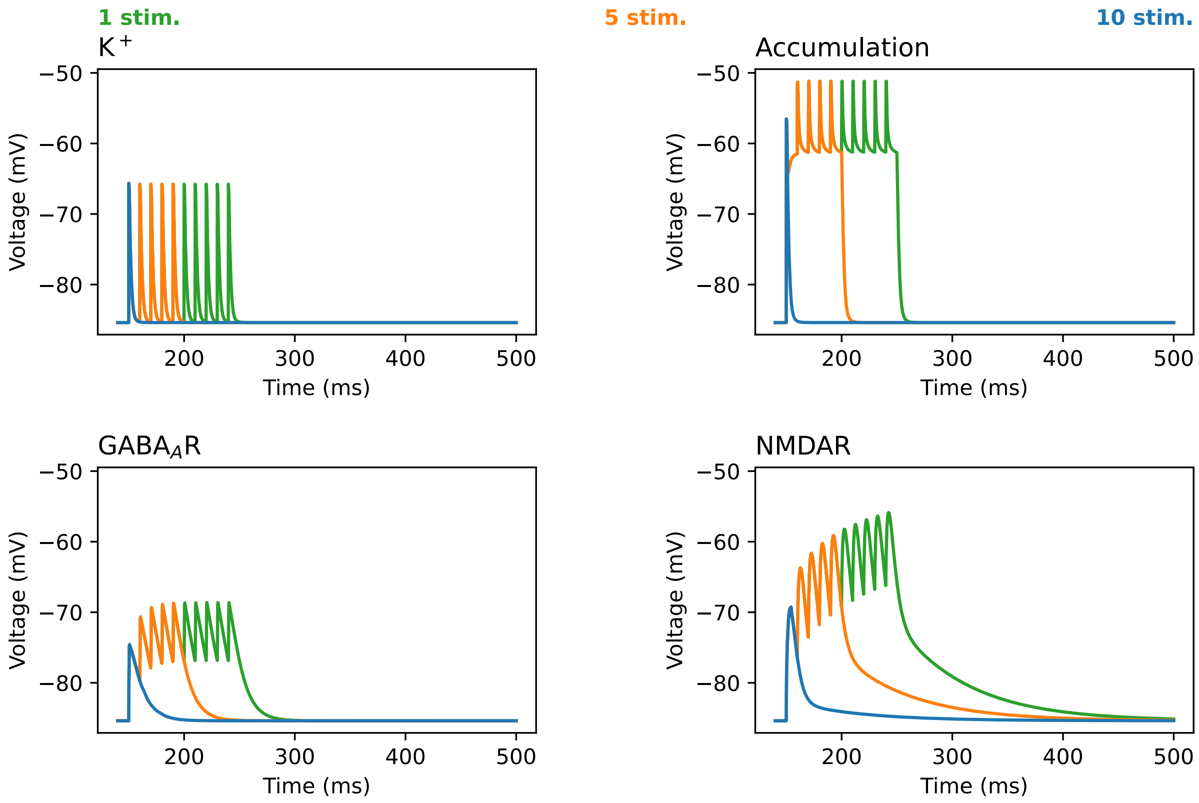


**Fig S10.** Voltage traces for experimental fits
Voltage traces of the model, when fitting to experimental results observing depolarization [24]. Four models are shown, Top-left) all $K^+$ channels ($K^+$; complements Fig 3 D), Top-right) all $K^+$ channels and slowing of $K^+$ dynamics (Accumulation; complements Fig 4 E), Bottom-left) all GABA receptors activated ($GABA_AR$; complements Fig 6 E), and Bottom-right) all glutamate-activated channels (NMDAR; complements Fig 7 E). The y-axis shows the membrane potential changes ($V_m$) within the Stimulated PAP for each model. Stimuli protocols are colored with the color code at the top of the figure.

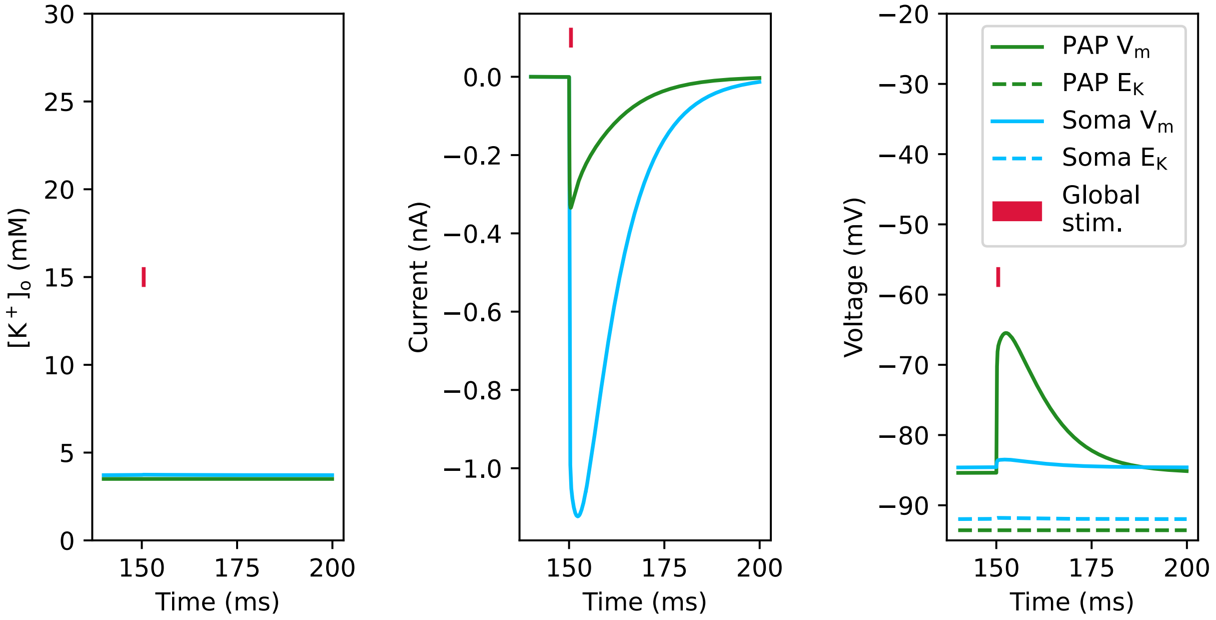


**Fig S11.** GABA response of hippocampal astrocyte caused by near soma puff
Changes in Left) [K$^+$]$_o$, Middle) currents, and Right) membrane potential for both soma and single PAP during the global 1.5 mM transient GABA puff application. The model considers a hippocampal astrocyte, where GABA$_A$R expression is more abundant compared to cortical astrocytes. The protocol qualitatively matches experimental protocols, and results from Chung et al. [52]. 440 GABA$_A$R are placed per model section to roughly match PAP densities in control conditions used in previous simulations.

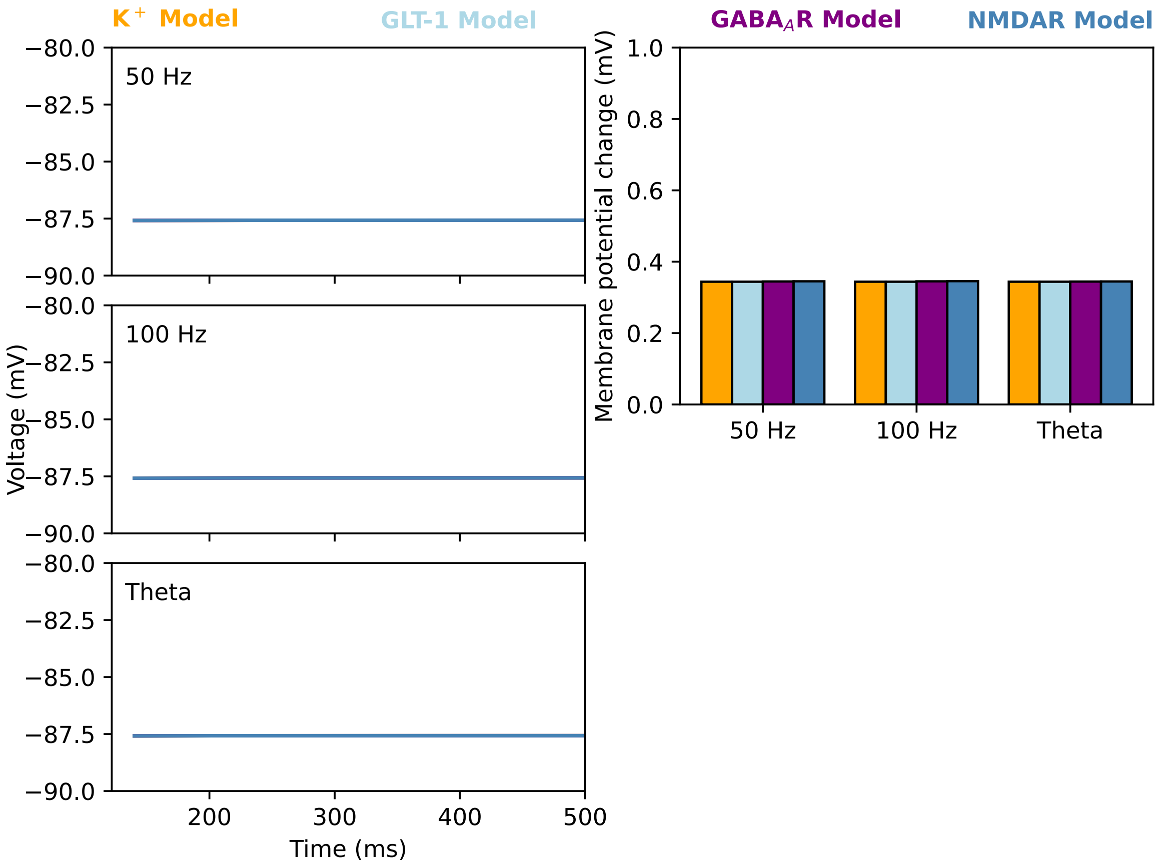


**Fig S12.** Somatic responses of the model to simultaneous synaptic activation
Responses to the combined stimulation of 40 PAPs when stimulating $K^+$, GLT-1, NMDAR, or GABA$_A$R models. The simulations are run with intracellular diffusion, and results were recorded at the soma during PAP stimulation. Left) Voltage traces at the soma during the 40 simultaneous stimulations of PAPs with each respective stimulation protocol. Right) Peak voltage amplitudes were measured at the soma during the simultaneous activation. There is no significant effect of simultaneous stimulation at the soma during these protocols.

### Supplementary Text

For all subsequent passages, $E_K$ was computed dynamically based on $[K^+]_o$. $T$ was the absolute temperature, which was constant and matched the experimental conditions of Armbruster et al. [24]. All neurotransmitter-related models were implemented as the NEURON point-process framework to ensure uniform stimulation and activation, see SI text S12.

Supplemental text S1: Derivation of $K^+$ diffusion flux in realistic extracellular space

Extracellular diffusion flux was computed as linear concentration-dependent steady-state flux from a radial extracellular shell compartment. Initially, the extracellular compartment was calculated as a shell with depth ($l_{\text{ecs}}$). Therefore, the extracellular hollow cylinder had an inner radius of ($r_a$) and outer radius of ($r_{\text{ecs}} = r_a + l_{\text{ecs}}$). Given an ion concentration defined by the astrocyte inner membrane, and the outside of the radial shell to be a constant bath concentration ($C_{\text{bath}}$), the change of ion concentration ($C(r)$) at radius ($r$) over time using Fick's second law of diffusion is,

$$\frac{dC(r)}{dt} = \frac{D}{r}\frac{d}{dr}\left(r\frac{dC(r)}{dr}\right) \tag{9}$$

where $D$ is the diffusion coefficient. Calculating the flux at steady-state yields,

$$C(r) = A\ln r + B \tag{10}$$

where $A, B$ are integration constants. Since at $t = 0$, $C(r_{\text{ecs}})$ is equal to the bath concentration, at the outer surface $r_{\text{ecs}}$, flux ($J_b$) is,

$$J_b = -\frac{D}{r_{\text{ecs}}}\frac{C(r_a) - C_{\text{bath}}}{\ln(r_a/r_{\text{ecs}})} \tag{11}$$

Since flux is in $\text{mol}\,\text{cm}^{-2}\,\text{s}^{-1}$, the total efflux ($J_{\text{tot}}$) out of the extracellular space, with effective membrane area ($A_{\text{eff}}$), extracellular space volume ($\text{Vol}_{\text{ECS}}$) and effective extracellular diffusion ($D_{\text{ECS}}$) is,

$$\begin{aligned} J_{\text{tot}} &= J_b * A_{\text{tot}}/\text{Vol}_{\text{ECS}} && (12)\\ &= J_b * \frac{2r_{\text{ecs}}\pi L}{\pi L(r_{\text{ecs}}^2 - r_a^2)} && (13)\\ &= -\frac{2 * D_{\text{eff}}}{\ln r_a/r_{\text{ecs}}(2l_{\text{ecs}}r_a + l_{\text{ecs}}^2)}(C(r_a) - C_{\text{bath}}) && (14) \end{aligned}$$

In this case, $D_{\text{eff}}$ is calculated as,

$$D_{\text{eff}} = \frac{D_{\text{K}}}{\tau_{\text{tor}}^2} \tag{15}$$

Parameters of depth can be found in Table 1 and Table S1.

#### Supplemental text S2: Kir 4.1 channel model

The Kir 4.1 channel model was based on Yim et al. [88]. It follows a conventional conductance-based model with a square-root dependence on $\text{K}_o^+$ [89]. Conductance was defined as,

$$g_{\text{Kir}} = \bar{g}_{\text{Kir}}\sqrt{[K^+]_o} \tag{16}$$

where $\bar{g}_{\text{Kir}}$ was the maximum conductance. $g_{\text{Kir}}$ was calibrated to fit experiments by Yang et al. [90], which were single channel conductance measured under 40 mM of $[\text{K}^+]_o$. $[\text{K}^+]$ in the astrocyte and the extracellular space gave the reversal potential ($\text{E}_\text{K}$). Additionally, the model contained one Hodgkin-Huxley (HH) type inactivation particle $l$, which fits Kir 4.1 recordings used in Stegen et al. [105].

$$l = (l_{inf} - l)/\tau_l \tag{17}$$

$l_{inf}$ and $\tau_l$ are given by,

$$l_{inf} = 1/(1 + exp((\text{V}_m - \text{V}_{halfl})/k_l)) \tag{18}$$

$$\tau_l = 1/((\tau_a exp(-\text{V}_m/\text{V}_{halft}) + \tau_b exp(\text{V}_m/\text{V}_{halft}))) \tag{19}$$

where $\text{V}_{halfl}$,$\text{V}_{halft}$ are the values of $l, t$ at the half of maximum, $\tau_a, \tau_b$ are time constant control parameters fit to experimental data. The model is defined below, with all parameter values mentioned in the Table S2.

$$I_{\text{Kir}} = g_{\text{Kir}} l(\text{V}_m - \text{E}_\text{K}) \tag{20}$$

#### Supplemental text S3: Channel number estimates

The channel numbers for Kir 4.1 and passive $\text{K}^+$ leak were fit to recreate the space constant of 3.6 µm at the soma (Fig S3), predicted by Zhou et al. [34]. The Kir 4.1 currents were marked as 50% of the necessary $\text{I}_\text{K}$, to match experimental results [22]. The passive $\text{K}^+$ leak conductance was determined as the residual current, excluding the effects of the other $\text{K}^+$ components (NKA).

#### Supplemental text S4: K2P TREK-1/TWIK-1 channel model

TWIK and TREK channels, which contribute to passivity in mature astrocytes, were modeled as a single passive ohmic channel with dependence on the $E_K$.

$$I_{K,L} = g_{K,L}(V_m - E_K) \tag{21}$$

where $g_{K,L}$ was the conductance and $I_{K,L}$ was the current of the leak channel. Parameters of conductance and reversal potential can be found in Table 1 and S3.

#### Supplemental text S5: Na/K-ATPase model

The Na/K-ATPase (NKA) model was based on the work of Somjen et al., which utilizes an analytical Hill-type reaction scheme for ion pumping [26]. The result was converted to current by utilizing a maximum current of 1 mA cm$^{-2}$ [26]. Specifically for the intracellular diffusion model, the model required a 150-fold increase in channel density to maintain resting membrane potential (RMP) for our sodium constant model. Therefore, the current was calculated as,

$$I_{NKA} = \bar{I}_{NKA} \frac{1}{(1 + \frac{[K^+]_o^b}{[K^+]_o})^2} \frac{1}{(1 + \frac{[Na^+]_i^b}{[Na^+]_i})^3} \tag{22}$$

Where $\bar{I}_{NKA}$ is the maximum current of the model, $[K^+]_o^b, [Na^+]_i^b$ is the baseline concentration for these ions, and $[K^+]_o, [Na^+]_i$ is the concentration of respective ions in those compartments. Parameters can be found in Table S4.

#### Supplemental text S6: NMDAR channel model

The current dynamics of the NMDAR were defined as follows, based on Moradi et al. [91].

$$I_{NMDAR} = g_{NMDAR}\ k_{Mg}(w_C C + w_B B - A)(V_m - E_{NMDAR}) \tag{23}$$

$$\frac{dA}{dt} = -\frac{A}{\tau_A} \tag{24}$$

$$\frac{dB}{dt} = -\frac{B}{\tau_B} \tag{25}$$

$$\frac{dC}{dt} = -\frac{C}{\tau_C} \tag{26}$$

The current of the NMDA channel ($I_{NMDAR}$) was modeled as a combination of three ordinary differential equations (ODEs) for $A, B, C$ with specific weights $w_B, w_C$ and time constants $\tau_A, \tau_B, \tau_C$. $A, B, C$ are initialized as 0, and upon activation, are increased by the synaptic factor ($f_{NMDA}$) calculated from time constants in the equation below.

$$f_{\text{NMDA}} = \frac{1}{-\exp(-t_{NMDA}/\tau_A) + w_B \exp(-t_{NMDA}/\tau_B) + w_C \exp(-t_{NMDA}/\tau_C)} \tag{27}$$

$$t_{NMDA} = \tau_A \tau_B \ln(\frac{\tau_B}{w_B \tau_A}))/(\tau_B - \tau_A) \tag{28}$$

$g_{\text{NMDAR}}$ and $\text{E}_{\text{NMDAR}}$ were the conductance and reversal potential, according to Ohmic Law.

The NMDA channel differs from neuronal channels due to its weak magnesium block, which was modeled as follows.

$$k_{\text{Mg}} = \frac{1}{1 + [\text{Mg}^{2+}]_O \ {K_0}^{-1} \ \exp(z \ \delta \ F \ (-\text{V}_m + s) \ R^{-1} \ T^{-1})} \tag{29}$$

Here, $[\text{Mg}^{2+}]_O$ was the extracellular concentration of magnesium. $z, F, R, T$ are thermodynamic constants corresponding to the valence of ions ($z = 2$ for magnesium), Faraday's constant, gas constant, and absolute temperature. $\delta$ is the relative electrical distance for the magnesium block. $\delta, s, K0$ were changed to match the weaker susceptibility of astrocytic NMDARs. $K_0$ is the $\text{IC}_{50}$ for extracellular magnesium at 0 mM. Parameters related to the weaker magnesium block were tuned to fit experimental results [12].

Channel conductance was dynamically determined by $A, B, C$ and by additional changes to the time constants, $\tau_A, \tau_B, \tau_C$, providing a HH-type gating scheme.

$$g_{\text{NMDAR}} = \bar{g}_{\text{NMDAR}} * (w_C C + w_B B - A) \tag{30}$$

$$\bar{g}_{\text{NMDAR}} = g_{VI} + g_{VD} \tag{31}$$

$$\frac{dg_{VD}}{dt} = (\text{V}_m - g_{VD0}) * g_{VDs} * g_{VI} \tag{32}$$

$$\tau_B = q_{10} \ (\tau_{B,0} + a_B \ \exp(\lambda_B \text{V}_m)) \tag{33}$$

$$\tau_C = q_{10} \ (\tau_{C,0} + a_C \ \exp(-\lambda_C \text{V}_m)) \tag{34}$$

$$q_{10} = Q_{10}^{(T_0 - T)/10} \tag{35}$$

where, $g_{VD}, g_{VD0}, g_{VI}, \tau_g$ were the voltage-dependent conductance, voltage shift factor, voltage independent conductance and time constant respectively. The equations were changed to qualitatively mimic the voltage dependence seen in Lalo et al. [12]. $g_{VD,0}$ was the final value of $g_{VD}$ when $t = \infty$. The voltage-independent component $g_{VI}$ was changed from 1 µS to 33 pS to match single channel recordings [56].

The dynamic time constants were defined as an exponentially rising constant with initial value $\tau_0$ and $a_i, \lambda_i$ obtained from experimental data fitting for respective time constants ($i$ =B,C). These time constants were multiplied by $q_{10}$, which converts these constants to temperature-dependent ones.

$T_0$ is the reference temperature for the given time constants measured experimentally. For desensitization, the original model utilized fast and slow dynamics. As repetitive stimulation protocols used in the current model were faster than those used in the experimental data, the fast desensitization factor was neglected. Slow desensitization occurred at 2500 ms time constant.

These equations summarize the three exponential model by Moradi et al. [91], values of parameters are shown below in Table S5.

Supplemental text S7: GABA$_A$R model

The GABA$_A$R model was a classic two-state synaptic conductance model adapted from Schulz et al. [94], with changes to the reversal potential to fit astrocytic conditions.

The GABA$_A$R two-state model equations,

$$I_{\text{GABAR}} = \bar{g}_{\text{GABAR}} * f(\text{V}_m) * (B - A) * (\text{V}_m - \text{E}_{\text{Cl}}) \tag{36}$$

$$f(\text{V}_m) = 1 + \frac{-0.75}{1 + \exp(\frac{\text{V}_m - \text{V}_{halfv}}{\text{V}_{halfl}})} \tag{37}$$

$$\frac{dA}{dt} = -\frac{A}{\tau_A} \tag{38}$$

$$\frac{dB}{dt} = -\frac{B}{\tau_B} \tag{39}$$

The current of the GABA$_A$R channel was modeled as a combination of two ODEs $A, B$. Each ODE had a time constant $\tau_A, \tau_B$. $A, B$ are originally initialized as 0, and upon activation, are increased by the synaptic factor ($f_{\text{GABA}}$) calculated from time constants in the equation below.

$$f_{\text{GABA}} = \frac{1}{-\exp(-t_{GABA}/\tau_A) + \exp(-t_{GABA}/\tau_B)} \tag{40}$$

$$t_{GABA} = \tau_A \tau_B \ln(\frac{\tau_B}{\tau_A}))/(\tau_B - \tau_A) \tag{41}$$

$f(\text{V}_m)$ was the voltage dependent rectification factor for GABA$_A$R, and was determined by $\text{V}_{halfv}$,$\text{V}_{halfl}$. $\bar{g}_{\text{GABAR}}$ and $\text{E}_{\text{Cl}}$ were the unitary conductance and chloride reversal potential, according to Ohmic Law. The main difference between the original model and our model was the change in reversal potential and unitary conductance, in order to match astrocyte dynamics. Parameters for each reaction rate are listed in the Table S6 and Table S7.

Supplemental text S8: Glutamate Transporter

The glutamate transporter model was modified from Savtchenko et al. [29], with slight alterations such as changing the MODL file from a membrane mechanism to a point process by adding a synaptic weight and calculating the membrane surface area within the MODL file. The model was a 6-state kinetic model with the reactions listed below. The glutamate dynamics were fit to a double-exponential equation, matching Savtchenko

| Reaction | Forward reaction rate | Backward reaction rate |
|---|---|---|
| $C1 \leftrightarrow C2$ | $[Glu]_o\ k12\ u(\mathrm{V}_m, -0.1)$ | $k21$ |
| $C2 \leftrightarrow C3$ | $[Na]_o\ k23\ u(\mathrm{V}_m, 0.5)$ | $k32$ |
| $C3 \leftrightarrow C4$ | $k34\ u(\mathrm{V}_m, 0.4)$ | $k43$ |
| $C4 \leftrightarrow C5$ | $k45$ | $k54\ [Glu]_i$ |
| $C5 \leftrightarrow C6$ | $k56u(\mathrm{V}_m, 0.6)$ | $k65\ [\mathrm{Na}^+]_i$ |
| $C6 \leftrightarrow C1$ | $k61\ [\mathrm{K}^+]_i$ | $k16\ u(\mathrm{V}_m, 0.6)\ [\mathrm{K}^+]_o$ |

et al. [29], shown below (Time constants Table S9).

$$[Glu] = \overline{[Glu]} \frac{\tau_2}{\tau_2 - \tau_1}(-\exp(-t/\tau_1) + \exp(-t/\tau_2)) \tag{42}$$

$t$ is the current time, $[Glu]$ is the glutamate concentration, and $\overline{[Glu]}$ is the maximal glutamate concentration determined by the stimulus input. $u(\mathrm{V}_m, k) = \exp(k\mathrm{V}_m/(53.4))$ for below kinetic rates.

where $\mathrm{V}_m, k$ were dependent on the aforementioned reactions. Current ($I_{\mathrm{GLT}}$) were calculated based on charges assigned to each reaction. Reaction between state $C1$ and $C6$ was designated for $\mathrm{I_K}$ and contributed to changes in $\mathrm{K}^+$. Parameters for each reaction rate are listed in Table S8.

Supplemental text S9: Sodium leak channels

Sodium leak channels followed conventional conductance-based equations.

$$I_{\mathrm{Na}} = g_{\mathrm{Na}}(\mathrm{V}_m - \mathrm{E}_{\mathrm{Na}}) \tag{43}$$

where $\mathrm{E}_{\mathrm{Na}}$ was the specific reversal potential calculated from the sodium Nernst potential. $g_{\mathrm{Na}}$ is the conductance, $\mathrm{V}_m$ was the membrane potential and $I$ was the current of the leak channel. The sodium conductance $g_{\mathrm{Na}}$ was calculated for each compartment to result in a RMP of -85 mV. As Kir 4.1 and $\mathrm{K}^+$ leak were the main components of hyperpolarizing current during RMP, the equation was,

$$g_{\mathrm{Na}} = (g_{\mathrm{Kir}} * l_{inf} * k_l + g_{\mathrm{K}})(\mathrm{RMP} - \mathrm{E_K})/(\mathrm{RMP} - \mathrm{E}_{\mathrm{Na}}) \tag{44}$$

$g_{\mathrm{Kir}}$ was single channel conductance for Kir 4.1, $l_{inf}$ was the value for the gating particle at rest at RMP. Parameters of conductance and reversal potential can be found in Table 1 and Table S2.

Supplemental text S10: Gap junction model

Gap junctions in the model were implemented as shunting passive currents independent of $\mathrm{K}^+$,

$$I_{\mathrm{gap}} = g_{\mathrm{gap}}(\mathrm{V}_m - \mathrm{V}_{\mathrm{adj}}) \tag{45}$$

Where $g_{\text{gap}}$ is the conductance, $V_m$ is the membrane potential, and $V_{adj}$ is the membrane potential of the adjacent astrocyte. Experimental values of unitary conductance measured in cultured astrocytes were implemented [106], with channel numbers fit to match trans-junctional conductances measured in brain slices [107]. The gap junctions were positioned only at the end of peripheral branches and $V_{\text{adj}}$ was equal to the RMP. Only under bath conditions were the theoretical adjacent astrocyte RMP values altered. The model was linear, as trans-junctional ranges tested in our model were within amplitudes that were not affected by gap junction closing [108]. Parameters can be found in Table S10.

#### Supplemental text S11: Derivation of analytical model for $K^+$ uptake extracellular space

To examine how changes in the ECS size would directly affect astrocyte uptake dynamics, we used an analytical model of $K^+$ uptake via Kir 4.1. The model considered 500 Kir 4.1 channels on a potential-clamped membrane for simplicity. We examined ECS sizes of depth 10 nm $\sim$ 100 nm which encompassed the ECS used in our model, and experimentally measured sizes [86]. The analytical model was integrated from the equations for Kir 4.1 currents (SI text S2) by converting current to ion flux. The resulting analytical model was,

$$[K^+]_o = \gamma \exp\left(2\text{Ei}^{-1}\left(-\frac{\beta t}{\sqrt{\gamma}\exp(\frac{\alpha}{2\beta})} + C\right) + \frac{\alpha}{\beta}\right) \tag{46}$$

$$\alpha = V_m \frac{\bar{g}_{\text{Kir}}}{zF\text{Vol}_{\text{ECS}}^{\frac{3}{2}}} \tag{47}$$

$$\beta = \alpha\frac{RT}{zF} \tag{48}$$

$$\gamma = \text{Vol}_{\text{ECS}}[K^+]_i \tag{49}$$

where, $\text{Ei}^{-1}$ is the inverse of the exponential integral, R,T,z,F are gas constant, temperature, valence, and Faraday's constant. $\bar{g}_{\text{Kir}}$, is the maximal Kir conductance and $\text{Vol}_{\text{ECS}}$ is the volume of the ECS. For initial conditions where $[K^+]_o$ is $[K^+]_o^{t=0}$ for $t = 0$,

$$C = \text{Ei}\left(\ln\left(\frac{\sqrt{[K^+]_o^{t=0}}}{\gamma} - \frac{\alpha}{2\beta}\right)\right) \tag{50}$$

#### Supplemental text S12: Conversion of synaptic weight to concentration

All values of synaptic weight in NEURON were converted to ligand concentration for our simulations. Synaptic weights were converted to concentration-dependent weights using the Hill-type binding equation below.

$$w = \frac{1}{1 + (\text{EC}_{50}/[L])^n} \tag{51}$$

Specific parameters are listed for NMDAR and $\text{GABA}_A$R in Table S11. Although GLT-1 was also implemented as a point process, this model used glutamate concentrations defined by exponential rise-decay dynamics with maximal amplitude matching synaptic weight to compensate for the kinetic nature of the model [29].

**Table S1.** Table for parameters of extracellular space model

| Parameter | Value | Units | Description | Comment/Ref. |
|---|---|---|---|---|
| $D_{\mathrm{K}}$ | $19.2 \times 10^{-6}$ | $\mathrm{cm^2\,s^{-1}}$ | Diffusion coefficient | [27] |
| $\tau_{\mathrm{tor}}$ | 1.6 | | Tortuosity | [27] |
| $l_{\mathrm{ecs}}$ | $40 \sim 100$ | nm | Depth of ECS compartment | [102] |

**Table S2.** Table for parameters of Kir 4.1 model

| Parameter | Value | Units | Description | Comment/Ref. |
|---|---|---|---|---|
| $\bar{g}_{\mathrm{Kir}}$ | 4.7 | pS | Maximum conductance for Kir | calibrated to match 50 pS at extracellular potassium concentration of 40 mM [90] |
| $V_{halfl}$ | -98.92 | mV | Voltage at half of maximum $l$ | Fit to data in Stegen et al. [105] |
| $k_l$ | 10.89 | mV | Sensitivity of $l$ | Same as above [105] |
| $\tau_a$ | 1.98 | $ms^{-1}$ | Control parameter for $\tau_l$ | [88] |
| $\tau_b$ | 1.44 $\times 10^{-2}$ | $ms^{-1}$ | Control parameter for $\tau_l$ | Same as above [88] |

**Table S3.** Table for parameters of TREK-1 model

| Parameter | Value | Units | Description | Comment/Ref. |
|---|---|---|---|---|
| $g_{\mathrm{K},L}$ | 1.15 | pS | Passive potassium conductance | calibrated to match space constant of 3.6 μm [22] |

**Table S4.** Table for parameters of Na/K-ATPase (NKA) model

| Parameter | Value | Units | Description | Comment/Ref. |
|---|---|---|---|---|
| $\bar{I}_{\mathrm{NKA}}$ | 1 | $\mathrm{mA\,cm^{-2}}$ | Maximal NKA current | [26] |

**Table S5.** Table of parameters for NMDAR model

| Parameter | Value | Unit | Description | Comment/Ref. |
|---|---|---|---|---|
| $[Mg^{2+}]_O$ | 1 | mM | Extracellular magnesium concentration | [101] |
| $K_0$ | 20 | mM | Half maximal concentration | Susceptibility to $Mg^+$ |
| $\delta$ | 0.1 | | Relative electrical distance | Susceptibility to $Mg^+$ |
| $s$ | 40 | mV | shifted $V_m$ dependence | Susceptibility to $Mg^+$ |
| $g_{VI}$ | 33 | pS | Maximal voltage independent conductance | [56] |
| $E_{NMDAR}$ | -0.7 | mV | Reversal potential | [101] |
| $g_{VDs}$ | 0.007 | $mV^{-1}$ | The slope factor for voltage dependent conductance | [109] |
| $g_{VD0}$ | -100 | mV | The $V_m$ at which $g_{VD,\infty}$ is zero | [109] |
| $w_B$ | 0.95 | | Percentage of decay for B | |
| $w_C$ | $1 - w_B$ | | Percentage of decay for C | |
| $a_B$ | 0.7 | ms | time constant parameter | [91] |
| $\lambda_B$ | 0.0243 | $mV^{-1}$ | time constant parameter | [91] |
| $a_C$ | 34.69 | ms | time constant parameter | [91] |
| $\lambda_C$ | 0.01 | $mV^{-1}$ | time constant parameter | [91] |
| $T_0$ | 26 | °C | reference temperature for Q10 | [91] |

| Time Constant | $T_0$ (°C) | $Q_{10}$ | Description | Comment/Ref. |
|---|---|---|---|---|
| $\tau_A$ | 1.69 | $2.2 \pm 0.5$ | Rising component | [110] |
| $\tau_{B,0}$ | 3.97 | 3.68 | Fast decaying component | [111] |
| $\tau_{C,0}$ | 41.62 | 2.65 | Slow decaying component | [111] |
| $\tau_g$ | 7 | 1.52 | Conductance change | [112] |

**Table S6.** Table of parameters for GABA$_A$R model

| Parameter | Value | Unit | Description | Comment/Ref. |
|---|---|---|---|---|
| $\bar{g}_{\mathrm{GABAR}}$ | 28 | pS | Unitary conductance | [57] |
| $V_{halfv}$ | -52 | mV | Voltage for half maximal rectification | [94] |
| $V_{halfl}$ | 3 | mV | Steepness of the rectification factor | [94] |
| $E_{\mathrm{Cl}}$ | -40 | mV | Chloride reversal potential | calculated from Nernst equation for model chloride conc. |

**Table S7.** Time constants for GABA$_A$R model

| Time Constant | Value | Unit | Comment/Ref. |
|---|---|---|---|
| $\tau_A$ | 0.1 | ms | [94] |
| $\tau_B$ | 10 | ms | [94] |

**Table S8.** Kinetic reaction rates for GLT-1

| Parameter | Value | Unit | Description | Comment/Ref. |
|---|---|---|---|---|
| $k12$ | 20 | $mM^{-1}\,ms^{-1}$ | Reaction rate from state 1 to 2 | [113] |
| $k21$ | 0.1 | $ms^{-1}$ | Reaction rate from state 2 to 1 | [113] |
| $k23$ | 0.015 | $mM^{-1}\,ms^{-1}$ | Reaction rate from state 2 to 3 | [113] |
| $k32$ | 0.5 | $ms^{-1}$ | Reaction rate from state 3 to 2 | [113] |
| $k34$ | 0.2 | $ms^{-1}$ | Reaction rate from state 3 to 4 | [113] |
| $k43$ | 0.6 | $ms^{-1}$ | Reaction rate from state 4 to 3 | [113] |
| $k45$ | 4 | $ms^{-1}$ | Reaction rate from state 4 to 5 | [113] |
| $k54$ | 10 | $mM^{-1}\,ms^{-1}$ | Reaction rate from state 5 to 4 | [113] |
| $k56$ | 1 | $ms^{-1}$ | Reaction rate from state 5 to 6 | [113] |
| $k65$ | 0.1 | $mM^{-1}\,ms^{-1}$ | Reaction rate from state 6 to 5 | [113] |
| $k16$ | 0.0016 | $mM^{-1}\,ms^{-1}$ | Reaction rate from state 1 to 6 | [113] |
| $k61$ | 2e-4 | $mM^{-1}\,ms^{-1}$ | Reaction rate from state 6 to 1 | [113] |

**Table S9.** Time constants for GLT-1 model

| Time Constant | Value | Unit | Comment/Ref. |
|---|---|---|---|
| $\tau_1$ | 0.61 | ms | [29] |
| $\tau_2$ | 5.8 | ms | [29] |

**Table S10.** Table for parameters of gap junction model

| Parameter | Value | Units | Description | Comment/Ref. |
|---|---|---|---|---|
| $g_{\text{gap}}$ | 56 | pS | Passive unitary conductance | Experimental measurement [106] |

**Table S11.** Table for parameters of Hill-based binding

| Parameter | Value | Unit | Comment/Ref. |
|---|---|---|---|
| NMDAR | | | |
| n | 1.2 | | [114] |
| $EC_{50}$ | 4.3 | µM | same as above |
| GABAR | | | |
| n | 1.5 | | [115] |
| $EC_{50}$ | 7 | µM | same as above |
